# Multi-step vs. single-step resistance evolution under different drugs, pharmacokinetics and treatment regimens

**DOI:** 10.1101/2020.10.19.344960

**Authors:** Claudia Igler, Jens Rolff, Roland R. Regoes

## Abstract

The success of antimicrobial treatment is threatened by the evolution of drug resistance. Population genetic models are an important tool in mitigating that threat. However, most such models consider resistance emergence via a single mutational step. Here, we assembled experimental evidence that drug resistance evolution follows two patterns: i) a single mutation, which provides a large MIC increase, or ii) multiple mutations, each conferring a small increase, which combine to yield high-level resistance. Using stochastic modeling we then investigated the consequences of these two patterns for treatment failure and population diversity under various treatments. We find that resistance evolution is substantially limited if more than two mutations are required and that the most efficacious drug type depends on the pharmacokinetic profile. Further, we demonstrate that, for resistance evolution in multiple steps, adaptive treatment, which only suppresses the bacterial population, is favored over aggressive treatment, which aims at eradication.

## Introduction

The rapid rise and spread of antimicrobial resistance severely curb the efficacy of drug treatments against pathogen infections. The choice of treatment strategy can significantly determine the efficacy of pathogen removal and the potential for resistance evolution^1–3^, highlighting the importance of careful consideration of drug type, dose and duration^1,4^. In order to deter drug resistance and preserve drug efficacy, treatment strategies should be guided by a predictive understanding of resistance evolution dynamics^5,6^ – a task that is substantially facilitated through mathematical modeling^1,5–7^. One severely understudied aspect in such approaches is that there are two fundamentally different patterns of *de novo* antibiotic resistance evolution^8^: i) ‘single-step’ resistance: a single mutation provides higher drug resistance than a given treatment dose, or ii) ‘multi-step’ resistance: the accumulation of several mutations of low individual benefit is necessary for high-level resistance. The availability of either pathway to a pathogen population under drug selection will affect the potential for resistance evolution and therefore the evolutionary dynamics in response to various treatment strategies.

The main class of models used to predict drug action are pharmacokinetic and pharmacodynamic (PKPD) models^7–10^, which describe the change in drug concentration over time (pharmacokinetics) and the corresponding effect on a pathogen population (pharmacodynamics). PKPD approaches are most commonly employed to study the efficacy of treatment without considering the possibility of resistance evolution, but coupled with bacterial population models, they can be used to investigate drug resistance evolution over time^11^. The minority of PKPD studies that do consider resistance evolution mostly assume that resistance evolves via a single, high-benefit mutation^1,8,11^.

In this study, we are going to comprehensively study the influence of the mechanistic pattern of resistance evolution itself (namely the benefits and costs of mutations) by considering ‘single-step’ resistance vs. ‘multi-step’ resistance. The emergence of mutations and their selection depend on an interplay between various treatment factors like drug type, dose and treatment duration. These factors have been studied before to various extent in isolation^1,4^, although rarely how their interactions shape resistance evolution^12,13^. We will first establish the existence of single-step and multi-step resistance patterns by reviewing evidence in the experimental literature, and then use the obtained parameter values to inform a stochastic PKPD model of multi-step resistance evolution, which we will explore under various treatment regimens.

We will establish the fundamental differences between evolutionary dynamics emerging from these two patterns in one specific treatment setting, but also explore the impact of various clinically relevant treatment strategies. First, we will compare two types of drugs, antibiotics (ABs) and antimicrobial peptides (AMPs). Antimicrobial peptides are key components of innate defenses but also important new antimicrobial drugs, which work by disrupting the bacterial membrane^14,15^ -as opposed to antibiotics, which generally target intracellular structures. AMPs have been found previously to significantly reduce the risk of resistance evolution compared to conventional antbibiotics^11,16^, partly explained by their distinct pharmacodynamics like higher killing rates^11^. Second, we will consider three different shapes of drug pharmacokinetics, which are all clinically relevant^1^, but have rarely been compared in a systematic manner^10,17^. These comprise fluctuating drug concentrations, increasing concentrations, which are then maintained at the highest level and finally constant, which can be achieved in high-dose IV interventions. Third, as a number of recent studies have questioned the practice of ‘radical pathogen elimination’^6,18,19^, we will compare aggressive elimination treatment with adaptive suppression - a strategy where the drug concentration is regularly adapted to the pathogen load - in a multi-step mutational framework^18,20^. Lastly, depending on the drug type, resistance evolution can be shaped either by chromosomal mutations or horizontal gene transfer (HGT) or both^21,22^. Assuming a scenario where both options are available, we will study the relative importance of resistance resulting from *de novo* mutations as compared to HGT, which plays an important role in antibiotic resistance evolution^21^, although likely not as much in antimicrobial peptide resistance^23^. Taken together, this will allow us to provide an empirically informed modeling framework, which predicts evolutionary dynamics of ‘single-step’ resistance vs ‘multi-step’ resistance in the context of drug type, pharmacokinetics and treatment strategies. Such frameworks provide critical tools for comparing the risk of drug treatment failure in clinical settings.

## Results

### Antibiotic resistance evolves via multiple low- or single high-benefit mutation(s)

Experimental studies document single target mutations as well as a sequence of mutational steps to drug resistance evolution in bacterial populations^24–29^, but no systematic review of these patterns has been conducted so far. Here, we only selected studies that report on both parameters, benefit and costs of resistance, ^24–29^ (see also methods) in order to obtain a complete picture of the mutational effects. We define the benefit and cost of a mutation as an increase in the minimum inhibitory concentration (MIC) and as a reduction in growth (in the absence of drug) respectively. Despite differences in study setup and type of resistance mutations, we clearly found a wide range of effects, with a large number of benefits below typical clinical MIC breakpoint values, which are often 10xMIC or higher^30^ (Table 1, Fig. 1) - hence likely necessitating multiple mutations for high resistance. The corresponding fitness costs range from almost none to a 25% reduction of the population growth rate and show a very weak positive correlation (R^2^=0.07, P=0.09) with (log) benefit over all studies taken together (Fig. 1B, S1). In general, mutations seem likely to incur more costs than benefits. Notably, our literature search suggests a difference in mutational benefit available for two different antimicrobials: the average benefit of resistance mutations to AMPs is substantially lower than for commonly used ABs (Table 1, Fig. S1). In the following, we use the correlation observed with these assembled benefit and cost values to inform a PKPD model that reflects the two patterns of resistance evolution.

**TABLE 1.**
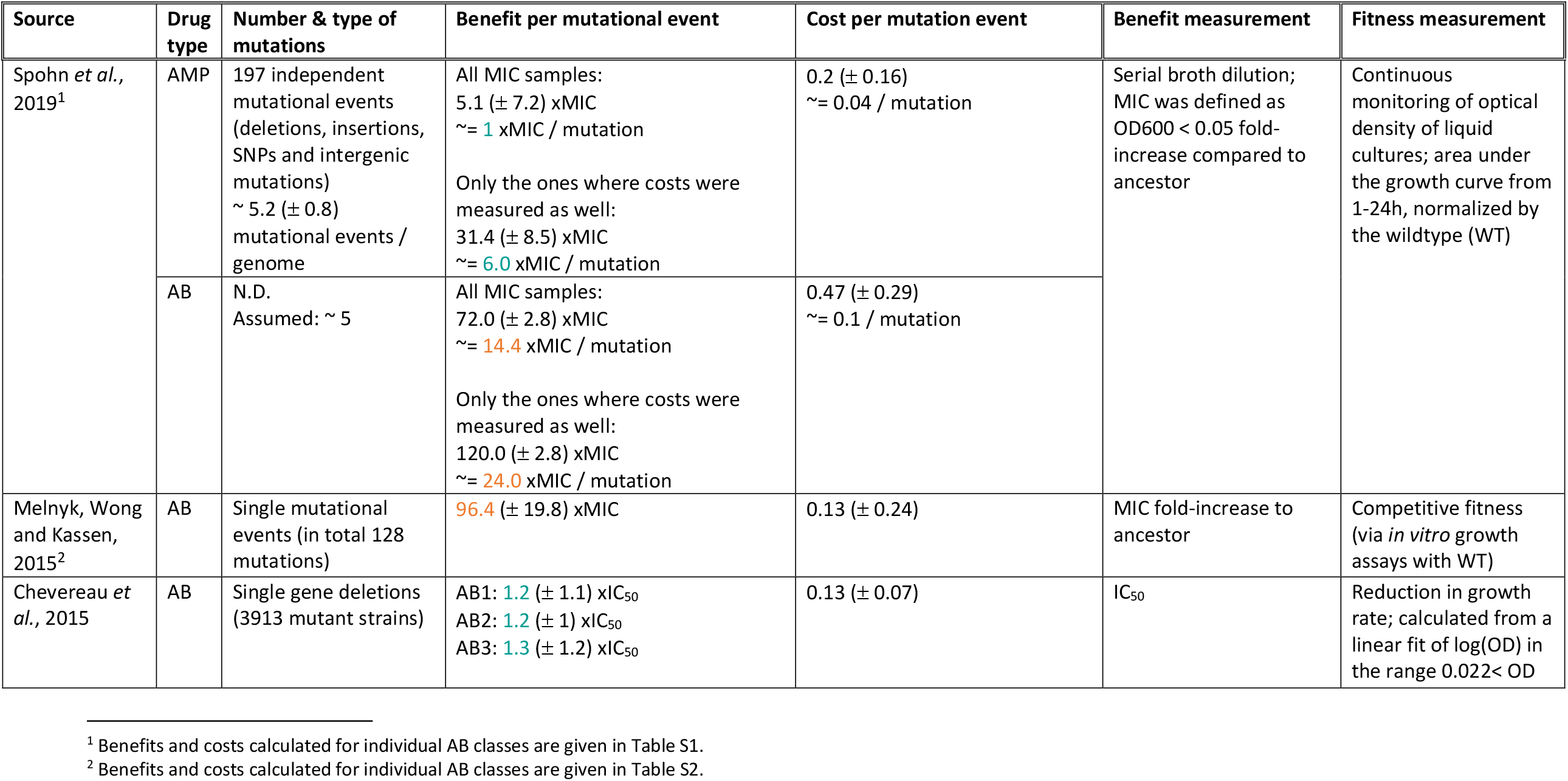
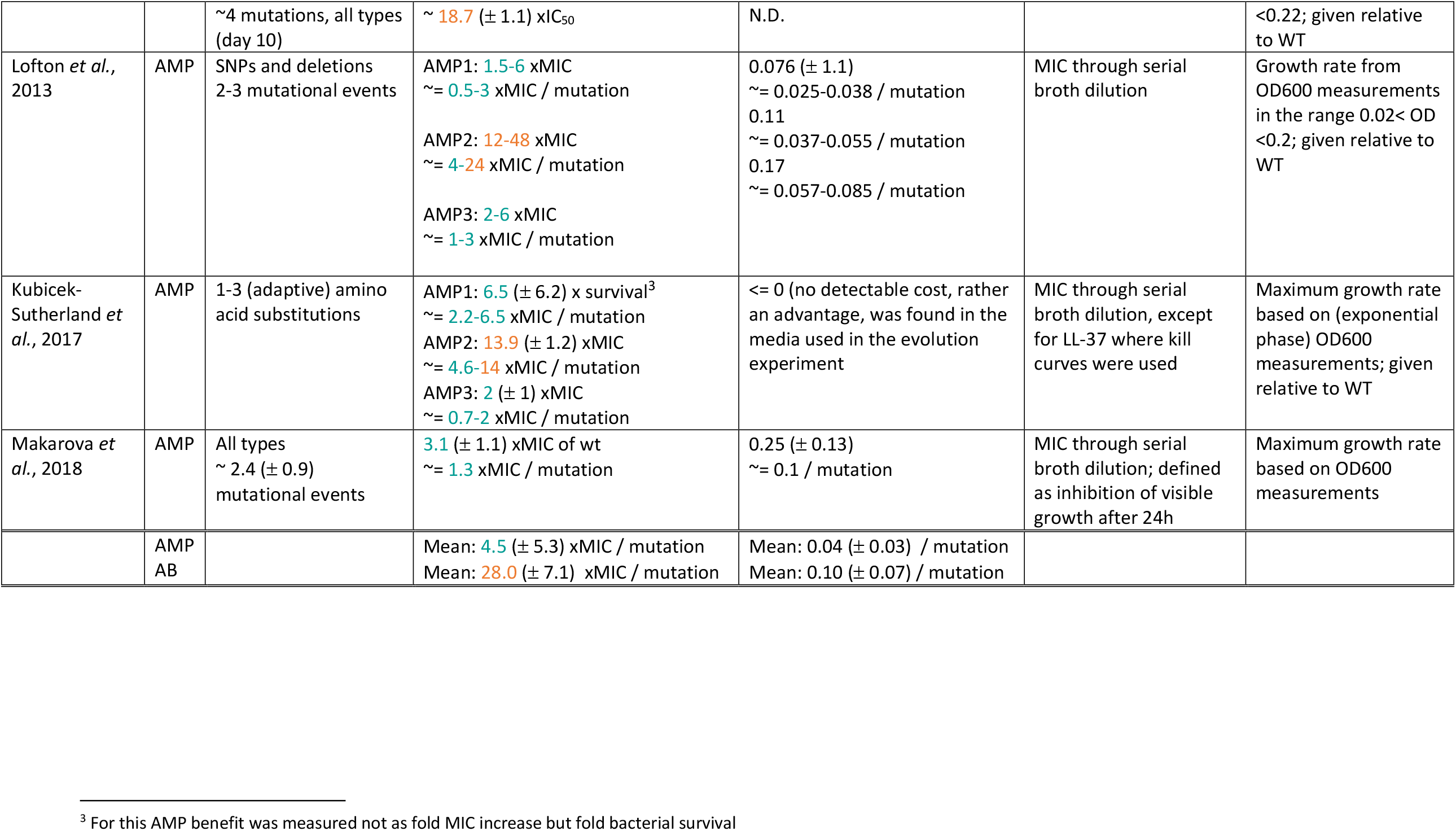
Benefits and costs of drug resistance mutations from experimental studies reported for antibiotics (ABs) and antimicrobial peptides (AMPs), with small mutational benefits (likely giving rise to multi-step resistance patterns) given in blue and large ones (likely giving rise to single-step resistance patterns) in orange, assuming a typical clinical drug dose of about 10xMIC (Fig. 1A,B).

| Source | Drug type | Number & type of mutations | Benefit per mutational event | Cost per mutation event | Benefit measurement | Fitness measurement |
| --- | --- | --- | --- | --- | --- | --- |
| Spohn *et al.*, 2019<sup>1</sup> | AMP | 197 independent mutational events (deletions, insertions, SNPs and intergenic mutations)<br>~ 5.2 (± 0.8) mutational events / genome | All MIC samples:<br>5.1 (± 7.2) xMIC<br>~ = 1 xMIC / mutation<br>Only the ones where costs were measured as well:<br>31.4 (± 8.5) xMIC<br>~ = 6.0 xMIC / mutation | 0.2 (± 0.16)<br>~ = 0.04 / mutation | Serial broth dilution; MIC was defined as OD600 < 0.05 fold-increase compared to ancestor | Continuous monitoring of optical density of liquid cultures; area under the growth curve from 1-24h, normalized by the wildtype (WT) |
|  | AB | N.D.<br>Assumed: ~ 5 | All MIC samples:<br>72.0 (± 2.8) xMIC<br>~ = 14.4 xMIC / mutation<br>Only the ones where costs were measured as well:<br>120.0 (± 2.8) xMIC<br>~ = 24.0 xMIC / mutation | 0.47 (± 0.29)<br>~ = 0.1 / mutation |  |  |
| Melnyk, Wong and Kassen, 2015<sup>2</sup> | AB | Single mutational events (in total 128 mutations) | 96.4 (± 19.8) xMIC | 0.13 (± 0.24) | MIC fold-increase to ancestor | Competitive fitness (via *in vitro* growth assays with WT) |
| Chevereau *et al.*, 2015 | AB | Single gene deletions (3913 mutant strains) | AB1: 1.2 (± 1.1) xIC<sub>50</sub><br>AB2: 1.2 (± 1) xIC<sub>50</sub><br>AB3: 1.3 (± 1.2) xIC<sub>50</sub> | 0.13 (± 0.07) | IC<sub>50</sub> | Reduction in growth rate; calculated from a linear fit of log(OD) in the range 0.022 < OD |

<sup>1</sup> Benefits and costs calculated for individual AB classes are given in Table S1.
<sup>2</sup> Benefits and costs calculated for individual AB classes are given in Table S2.

|  |  |  |  |  |  |  |
| --- | --- | --- | --- | --- | --- | --- |
| | | ~4 mutations, all types (day 10) | ~ **18.7** ($\pm 1.1$) xIC<sub>50</sub> | N.D. | | <0.22; given relative to WT |
| Lofton *et al.*, 2013 | AMP | SNPs and deletions<br>2-3 mutational events | AMP1: **1.5-6** xMIC<br>~ = **0.5-3** xMIC / mutation<br>AMP2: **12-48** xMIC<br>~ = **4-24** xMIC / mutation<br>AMP3: **2-6** xMIC<br>~ = **1-3** xMIC / mutation | 0.076 ($\pm 1.1$)<br>~ = 0.025-0.038 / mutation<br>0.11<br>~ = 0.037-0.055 / mutation<br>0.17<br>~ = 0.057-0.085 / mutation | MIC through serial broth dilution | Growth rate from OD600 measurements in the range 0.02 < OD < 0.2; given relative to WT |
| Kubicek-Sutherland *et al.*, 2017 | AMP | 1-3 (adaptive) amino acid substitutions | AMP1: **6.5** ($\pm 6.2$) x survival<sup>3</sup><br>~ = **2.2-6.5** xMIC / mutation<br>AMP2: **13.9** ($\pm 1.2$) xMIC<br>~ = **4.6-14** xMIC / mutation<br>AMP3: **2** ($\pm 1$) xMIC<br>~ = **0.7-2** xMIC / mutation | <= 0 (no detectable cost, rather an advantage, was found in the media used in the evolution experiment) | MIC through serial broth dilution, except for LL-37 where kill curves were used | Maximum growth rate based on (exponential phase) OD600 measurements; given relative to WT |
| Makarova *et al.*, 2018 | AMP | All types<br>~ 2.4 ($\pm 0.9$) mutational events | **3.1** ($\pm 1.1$) xMIC of wt<br>~ = **1.3** xMIC / mutation | 0.25 ($\pm 0.13$)<br>~ = 0.1 / mutation | MIC through serial broth dilution; defined as inhibition of visible growth after 24h | Maximum growth rate based on OD600 measurements |
| | AMP<br>AB | | Mean: **4.5** ($\pm 5.3$) xMIC / mutation<br>Mean: **28.0** ($\pm 7.1$) xMIC / mutation | Mean: 0.04 ($\pm 0.03$) / mutation<br>Mean: 0.10 ($\pm 0.07$) / mutation | | |

<sup>3</sup> For this AMP benefit was measured not as fold MIC increase but fold bacterial survival

**Figure 1.**
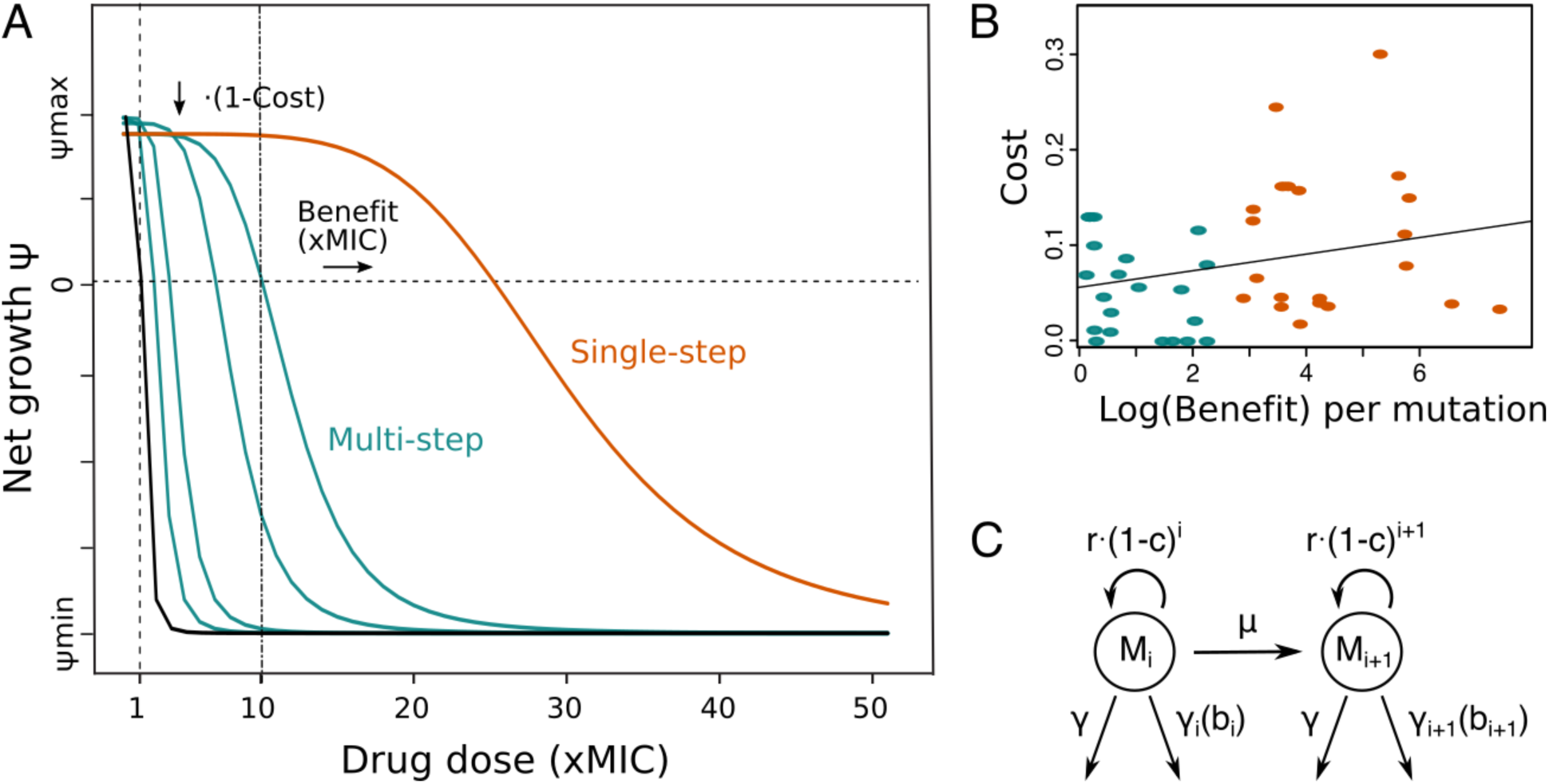
Pharmacodynamic (PD) model of single- and multi-step resistance. A) The PD curve relating bacterial net growth Ψ (which is between the maximal growth rate *ψ*_*min*_ and the maximal killing rate *ψ*_*min*_) with antimicrobial drug concentration (given in fold MIC) illustrating a sensitive wildtype (black) and mutants with either small (blue) or big (orange) MIC increases per mutation (benefit), assuming a typical clinial drug dose of 10xMIC. B) Shown are costs and benefits from various studies (Table1), each dot representing resistance mutations to a specific AB or AMP class. The cost of a mutation shows a very weak positive correlated with the log (benefit) (R^2^=0.07, P=0.09). Blue and orange colors show multi- or single-step resistance benefits given the drug dose in (A). C) Schematic of the PD model with several mutated subpopulations (*M*_*i*_), which grow with a cost r(1-c)^i^, determined by the number of mutations i, mutate with rate µ, and die at a constant rate *γ* and a drug-specific rate *γ*_*i*_(*b*_*i*_)).

### The pharmacodynamic model

We investigated the effect of an antimicrobial drug on growth and resistance evolution in a pathogen population over time by extending a previously described stochastic PKPD model (Methods)^11^. We considered however not only a single resistance mutation, but the potential emergence of a sequence of mutations, with each mutation conferring a certain (additional) benefit and cost (Fig. 1). The number of mutations needed for ‘full’ resistance depends on the applied drug dose, but generally low mutational benefits are more likely to necessitate multi-step resistance evolution. To compare scenarios where a single mutation is sufficient to scenarios where several mutations have to arise in one cell, we ran the simulations over a range of mutational benefits (2-100xMIC) – and their correlated fitness costs (Table 1, Fig. 1) – in combination with various drug doses (0.5-100xMIC). Hence, the minimum number of mutations necessary for resistance was pre-determined (Fig. S2), and we investigated how this affects the potential for pathogen survival and mutational diversity under various treatment strategies (PKs) and for two different antimicrobials (PDs) as described below. Competition between the subpopulations was modeled via imposing a carrying capacity for bacterial growth and very low turnover as soon as this capacity is reached.

### Multi-step resistance patterns show lower risk of treatment failure and lower genetic diversity

First, we determined the probability of treatment failure by simulating change of the pathogen population over 200 hours under treatment with drugs (PD parameters typical for ABs^11^) being applied once every 24h (peak pharmacokinetics). We assumed that the pathogen population initially consists of completely susceptible bacteria. We found that treatment failure (pathogen survival) was always close to 1 for single-step resistance evolution, but decreased rapidly if multiple mutations are required. Notably, already if 3 mutations were necessary to overcome the applied dose, the probability of pathogen survival approached 0 (Fig. 2A,S2). The qualitative picture of these results was not dependent on the specific cost-benefit correlation that we are assuming for most of our simulations (Fig. S3).

**Figure 2.**
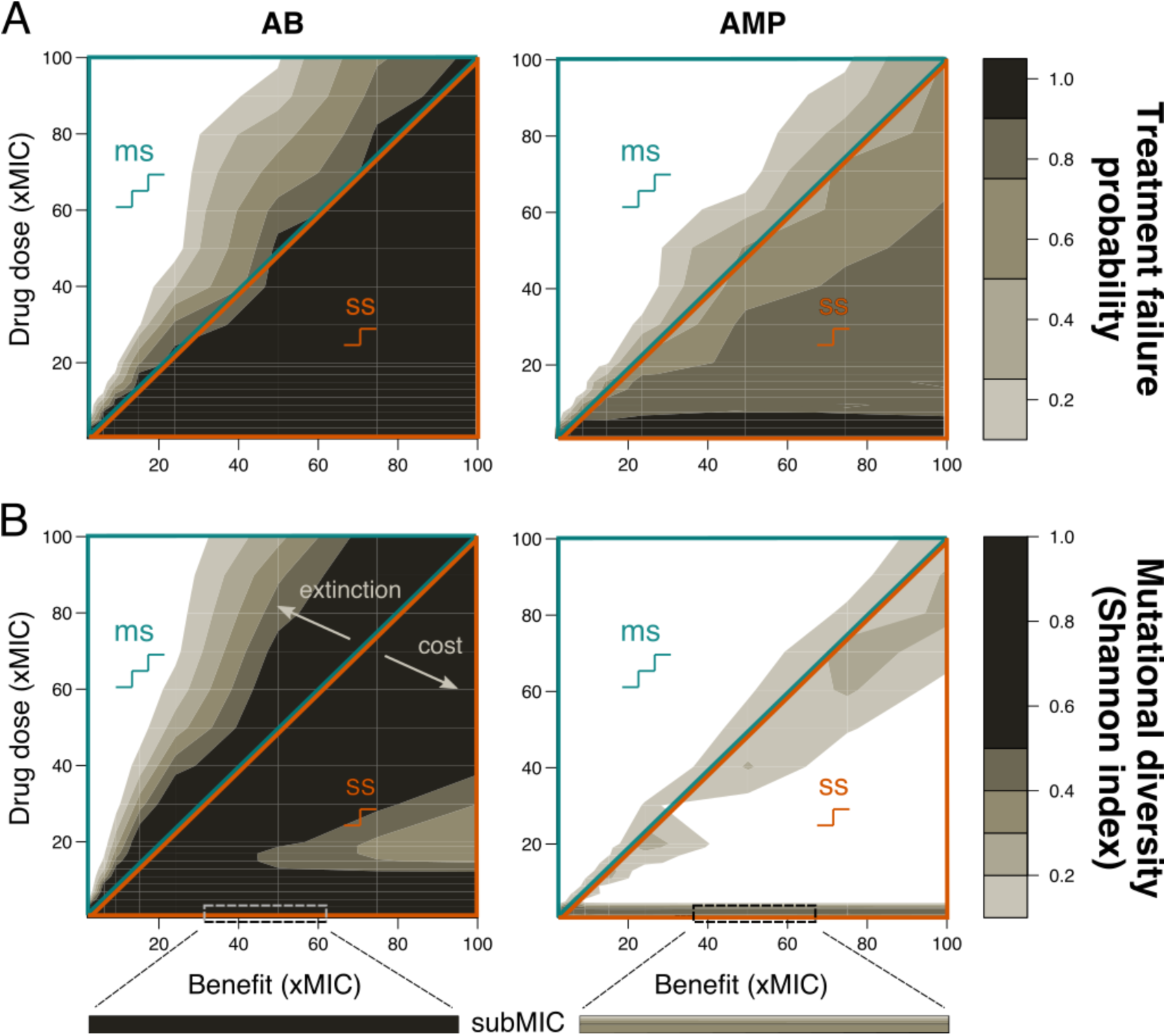
Resistance evolution with single- and multi-step patterns for peak PK. A) Treatment failure probability and B) mutational diversity are shown for two different antimicrobial classes (Abs – left, AMPs – right) for different combinations of mutational benefits (xMIC) and drug doses (xMIC). The diagonal line separates single-step (ss, lower orange triangle) from multi-step (ms, upper blue triangle) resistance. The arrows indicate the decrease in diversity either due to increasing extinction or due to increasing cost per mutation. A representative example of subMIC mutational diversity is shown magnified below the plots in B).

One aspect of resistance evolution that is especially important when considering several mutations is the mutational diversity that arises in the pathogen population: high genetic diversity (which we here assume is related to resistance) increases the probability that some individuals will be able to survive a given environment - such as treatment with other drugs - and increases the adaptive potential^31^. Using the Shannon index to determine the average mutational diversity in the population over the whole treatment period, we found a clear correlation between diversity and the resistance pattern: higher diversity attained with single-step resistance evolution. In particular, high diversity only occurs at low drug doses and when the mutational benefit is close to the applied dose (Fig. 2B), even if we increase the mutation rate proportionally to the number of mutations required (Fig. S4). It can be shown analytically that a mutant strain can invade at the mutant-free equilibrium if the death rate of the sensitive strain is higher than the death rate of the mutant normalized by the cost of the mutation(s) (Methods). The observed association between diversity and the resistance pattern is counter-intuitive(Fig. 2B), but can be explained as follows: at high drug doses and low benefits, this effect is due to extinction that effectively reduces genetic diversity, while at low doses and high benefits, high mutational costs inhibit the build-up of diversity. These findings agree with an experimental study showing that resistance alleles with low costs are favored^32^.

### Consistently lower treatment failure with multi-step resistance for various PKs and PDs

Our results clearly show less resistance if multiple mutations are necessary, but the relative importance of the number of resistance mutations over other treatment considerations like the type of drug (PD)^11,24^ or the application mode (PK) are unclear. Hence, we compared three different PKs: ‘peak’ (fast absorption and exponential decay), ‘ramp’ (slow, linear absorption and no decay) and ‘constant’ (immediate absorption and no decay) (Fig. 3A) Whereas constant PKs distinctly lowered the probability of treatment failure and the emergence of mutational diversity, peak and ramp PKs showed similar magnitudes of resistance evolution (Fig. 3B,C,S5,S6). However, ramp PKs lead to more than twice the mutational diversity with multi-step resistance patterns (Fig. S5), which suggests that treatment failure and pathogen diversity are connected in a non-trivial manner: while higher mutational diversity increases the risk of resistance evolution, neither its presence, nor absence, are obviously predictive of the treatment outcome (Fig. 2,S5,S6).

**Figure 3.**
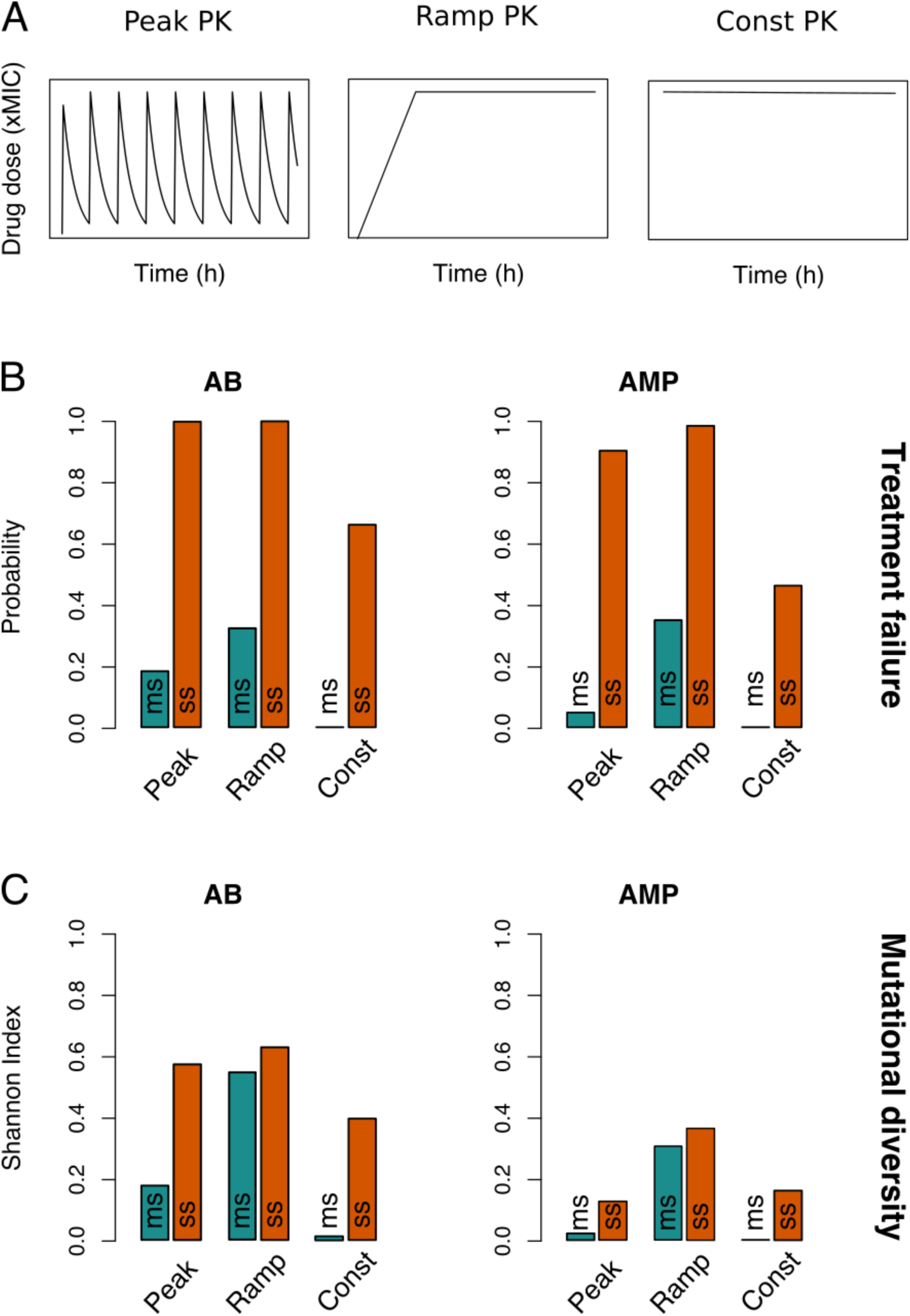
Resistance evolution patterns with different pharmacokinetics (PKs). A) The three PKs used in the model are shown over time (h) for the same peak drug concentration (xMIC). B) The treatment failure probabilities and C) Mutational diversities are given for the three PKs from (A) and two antimicrobial drug classes (ABs and AMPs). Blue (orange) bars show averages over the triangular multi-step (single-step) resistance areas in Fig. 2,S5,S6.

The evolutionary dynamics become even more complex if we consider different antimicrobial drugs, AMPs and ABs, by using two different PD parameter sets (Methods, Table S1). Briefly, AMPs have higher killing rates, steeper dose-response curves and lower mutation rates than ABs^11^. Consistent with previous findings that AMPs lead to a lower risk of resistance evolution and a narrower mutant selection window (MSW) than ABs ^11^, treatment failure and mutational diversity was lower for AMPs with peak and constant PK treatments (Fig. 2,3,S6). Notably, in accordance with other experimental studies^33^, we generally see mutations accumulating at sublethal drug doses, but the diversity is substantially lower in AMP treatments (Fig. 2,S3).

Interestingly, the steeper dose-response curve of AMPs seems to make their resistance dynamics more sensitive to the shape of the PK than those of ABs (Fig. 2,3,S5-S7): in contrast to the other two treatments, ramp PKs lead to a drastic increase in treatment failure with AMPs - especially in multi-step scenarios (Fig. 3, S5). Accordingly, for ramp PKs AMPs did not perform better and under some conditions even worse than ABs (Fig. S7). By varying the ramp duration (or equivalently the rate of drug uptake), we found that there is an intermediate range (48-84h), which showed increased treatment failure with AMPs over ABs (Fig. S8A). Paradoxically, while a narrow MSW generally hinders the emergence of numerous mutations in the population, for ramp PKs it leads to optimal selection conditions for the sequential emergence of increasingly higher resistance mutations due to the strong selection for the next mutation combined with sufficient time for its emergence. Hence, especially the risk of multi-step resistance is increased if AMPs are used with ramp treatments, as compared to the other PKs (Fig. 3B,C). The broader selection windows of ABs on the other hand overlap and mutations are less strongly favored (Fig. S8B). Overall, the number of resistance mutations was the main determinant of treatment outcome, but we also found a complex dependence on PK and PD characteristics.

### Multi-step resistance can lower the threshold for adaptive treatment application

So far we modeled the conventional treatment goal of ‘eradicating’ the pathogen population, but it has been suggested that under certain conditions ‘mitigation’ could be a superior strategy^18–20^, for example if it is likely that a resistant subpopulation already exists at the beginning of the treatment. This strategy is called adaptive treatment as drug doses are adapted to keep the sensitive population as big as possible and the total pathogen burden below a given limit (although in practice only total burden can be measured easily). The sensitive population provides a benefit by competitively inhibiting the resistant subpopulation, but also a risk by supplying mutational input (Fig. 4). This trade-off creates a threshold for the size of the pre-existent resistant subpopulation above which adaptive treatment type is preferable to aggressive ‘eradication’^18^.

**Figure 4.**
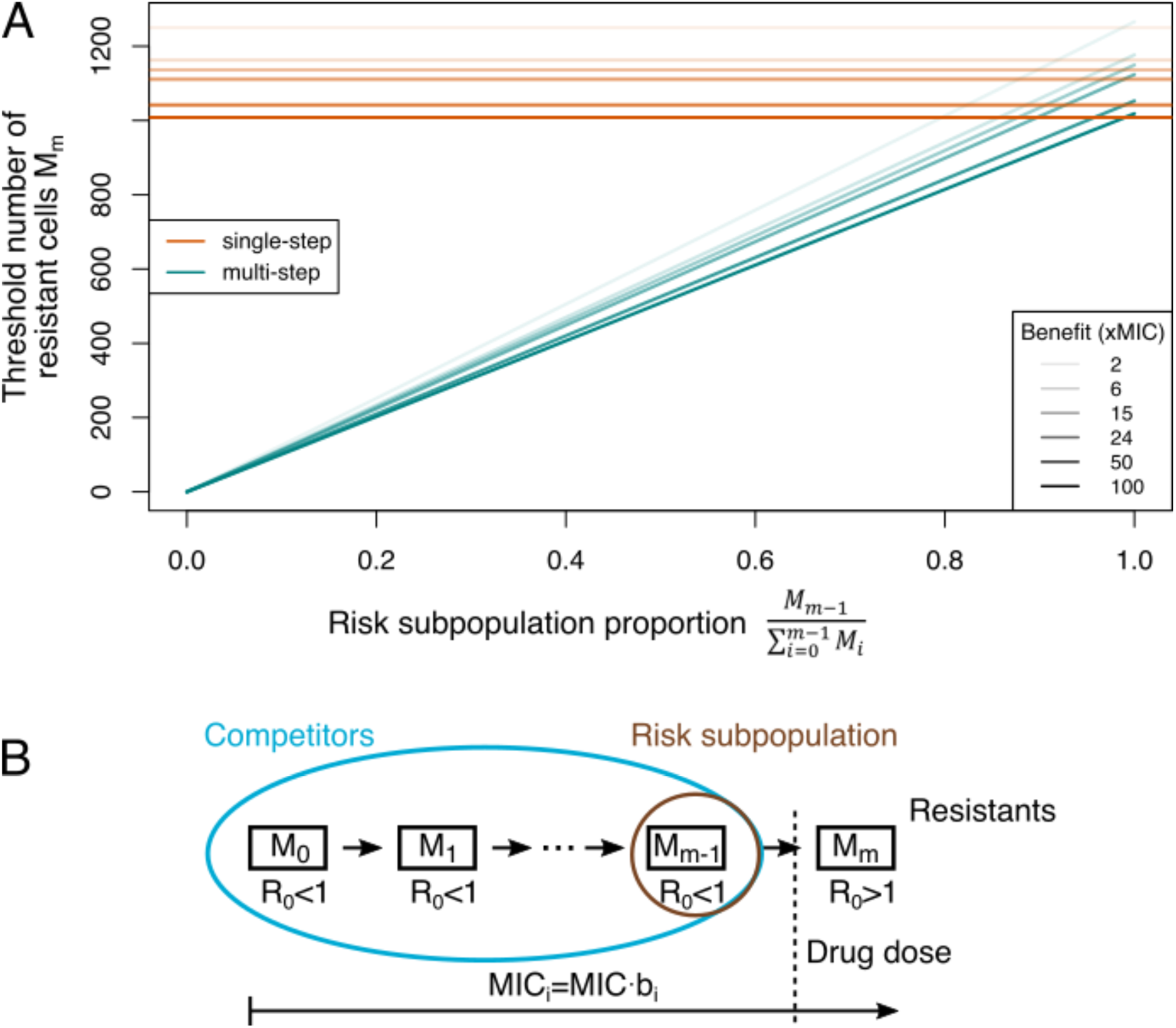
Adaptive treatment threshold. A) The dependence of the threshold number of resistant bacteria cells M_m_ is given for different proportions of the risk population to the whole competitor population for single-step (orange) and multi-step (blue) resistance patterns. Different benefits (and correlated costs) per mutation are shown as different line types. B) The MIC increases with every mutation (given by b_i_), but only an MIC above the given drug dose will lead to a reproductive number R_0_>1, i.e. growth of the population (resistant cells). All other subpopulations serve as competitors and the subpopulation one mutation away from resistance is the risk population.

Previously, the threshold for adaptive treatment was studied in a single-step resistance scenario^18^, but when we incorporated adaptive treatment in our multi-step resistance framework (Fig. S9), we found that the resistant subpopulation threshold above which adaptive treatment is more beneficial could potentially be much lower in the multi-step scenario than in the single-step framework (Fig. 4A). This can be intuitively explained by the fact that all (partially) sensitive bacteria serve as competitors for fully resistant cells, but only the subpopulation one mutation away from being fully resistant constitutes the risk population (Fig. 4B). Thus, with multi-step resistance there is a smaller population to supply resistant bacteria than with single-step resistance, rendering adaptive treatment more favorable. Additionally, (considering populations where adaptive treatment is favorable) the difference between adaptive and aggressive treatment in the duration until treatment failure can be several-fold larger for multi-step than single-step resistance patterns (Fig. S10). Hence, assuming either single- or multi-step evolution could lead to significantly different treatment strategy recommendations.

### Horizontal gene transfer does not change the treatment failure probability

In addition to chromosomal mutations^22^, antimicrobial resistance can be conferred through horizontal gene transfer^21^, which could facilitate resistance in multi-step scenarios. Therefore, we extended the model to allow for acquisition of a gene conferring full resistance at a low rate from the environment, and at a density-dependent rate from other cells carrying the HGT gene. The HGT gene always provided immediate resistance to the applied maximal dose, regardless of the benefit or costs conferred by mutations. We assumed that resistance through mutation or HGT can be acquired independently of each other and that their effects are multiplicative.

Even though HGT-carriers dominated the remaining pathogen population at the end of the treatment (Fig. S11), the addition of HGT did not change the probability of treatment failure (Fig. S12). This result holds true as long as the acquisition rate from the environment is lower than the mutation rate. Consequently, initial rescue of the population is due to mutations - and therefore dependent on the magnitude of the mutational benefit - whereas HGT resistance is acquired later during the infection, after which it spreads rapidly.

## Discussion

In this study we investigated the role of the drug resistance mechanism under various treatment designs by comparing multi-step to single-step evolution. We showed that these resistance patterns are relevant by gathering evidence of multi-step resistance patterns in the experimental literature (Table 1, Fig. 1). We then demonstrated that the number of mutations necessary for resistance significantly changes predictions for treatment outcome and optimality. Experimental support for our simulation results comes from studies reporting that mutational input limited to low benefits^8^ leads to decreased drug resistance evolution as compared to when high-benefit mutations are available^34^. Moreover, limited access to high-benefit mutations seems to curtail MIC increase beyond a certain threshold^25^.

The pattern of resistance evolution (single- and multi-step) is likely to be associated with the molecular mechanisms of resistance for a given antimicrobial: as an overall rule the magnitude of the resistance benefit correlates with the mechanism of resistance, e.g. efflux pumps yield low benefits, whereas specific drug target mutations yield high benefits^35^.

Unfortunately, the specific mutations linked to the benefit and cost of mutations in our literature analysis (Table 1) are generally not known. Overall however, MIC increase was low for drugs, which typically show unspecific resistance mechanisms via two-component systems or LPS modifications - as generally seen for AMP resistance^27–29^ -and high for drugs with typical resistance via specific target modifications, as seen for some AB classes (e.g. rifampicin resistance via RNAP subunit mutations)^36^. For most drugs the prevalent resistance mechanisms are known^21^, hence this information can be used to determine drug and dosing regimens that minimize resistance evolution based on the inferred pattern of resistance evolution (i.e. using the probability that a single- or multi-step pattern is underlying resistance evolution). A recent study also suggests that resistance evolution in biofilms is prone to occur through unspecific mechanisms (even if specific mechanisms are favored in planktonic cultures)^37^.

The risk of resistance evolution does not seem to be related to the underlying mutational diversity in the population in a trivial manner (Fig. 2,S5,S6). Reducing mutational diversity is however a worthwhile goal in its own right, as mutational diversity can increase adaptation by fixing more mildly deleterious mutations, which can then act as stepping stones for multi-drug resistance evolution^31^. We find that mutational diversity arises from a combination of selection pressure, bacterial growth and fitness costs and cannot be predicted from the mutational benefit or the probability of treatment failure alone. Further, diversity can be shaped in unexpected ways by interactions between the drug type and drug concentration changes, making drug choice not only dependent on the PD characteristics, but also the specific drug PK in the target body compartment. Notably, this can lead to recommendations of a specific drug application mode for one type of drug (e.g. AMPs for bolus drug application), but a different mode for another drug (e.g. ABs for drug infusions). The unexpected complexity in predicting optimal treatment strategies highlights the importance of questioning and extending assumptions such as single-step resistance made in current PKPD models. Only then will we be able to extract the drug characteristics crucial to reducing resistance evolution. This is exemplified by the steepness of the PD curve, κ: by analyzing the selection coefficients for various treatments, we find that κ governs not only the size of the MSW^11,25^, but generally shapes the selection pressure for resistance evolution in a qualitative manner. *ψ*_*min*_, the minimal bacterial growth rate, on the other hand, leads to substantial quantitative changes in selection pressure, meaning that κ and *ψ*_*min*_ shape the form and strength of drug selection independently (Fig. S8C). Ultimately, the interactions between PD and PK characteristics give rise to complex fitness landscapes that are navigated by mutations of various benefit and cost sizes.

Interestingly, AMP-like drugs show significantly more resistance evolution with ramp PKs than in the other PK scenarios. This is noteworthy as AMP expression patterns in the producing organisms resemble ramp PKs^38,39^. This finding could suggest another reason why natural AMP production in cocktails is favorable^40^, as AMP cocktails will limit the selection pressure and potential for resistance evolution to individual components. For clinical settings, our simulations caution that attention should be paid to the drug application mode when using AMPs. Indeed, AMP-like colistin and daptomycin are typically applied as (short) IV treatments^41,42^, which resemble peak PKs, and they are still active as last-resort drugs for multi-drug resistant bacterial pathogens^41,42^. Overall, our results agree with Yu *et al*., 2018 in that AMP treatment lowers resistance evolution and mutational diversity. This is particularly notable for multi-step evolution patterns, which so far seems to be the predominant mechanism how AMP resistance evolves (Table 1)^24,43–45^, thereby suggesting another advantage of AMPs over ABs, for which both evolution mechanisms are common^8,46–49^.

Unfortunately, distributions of mutational effects have rarely been characterized experimentally for drug resistance, and even then only for a single mutational step^25^. We show however, that this information is crucial as input for PKPD models to accurately predict resistance evolution and population diversity in response to drug treatment. Moreover, the empirical data that we used to inform our simulations did not provide explicit information about potential compensatory mutations, which arguably can influence the dynamics of resistance evolution^50^ – although likely in a very complex manner, as recent studies suggest^51^. According to our results, these mutations might even be a necessary means to allow multi-step resistance patterns to arise. If they emerge fast enough to compensate for the cost of the first mutation, they would increase the selection coefficient of this mutational subpopulations and thereby provide a stepping stone to high-level resistance. This might either be akin to crossing a fitness valley, if the first mutation does not provide a benefit, or it might facilitate climbing a fitness peak by making low-benefit mutations more favorable.

For many antimicrobial drugs resistance evolution cannot only arise through chromosomal mutations, but also by acquisition of resistance genes through HGT^21^. Notably, our results highlight the importance of transfer rates as we find rescue of the pathogen population through HGT resistance only if the initial acquisition rate is higher than the mutation rate. HGT is not only dependent on the recipient population size but also on the donor population size, hence using typical experimentally measured conjugation rates of 10^−11^-10^− 13^ ml cell^-1^h^-1 52^, environmental donors have to be more abundant than 10^5^ cells ml^-1^ to be faster than chromosomal mutation rates of 10^−6 53^, which might not always be the case at bacterial infection sites^54^. This implies either i) that HGT resistance is acquired after chromosomal mutations, ii) that HGT spreads mostly at sublethal drug doses, or iii) that acquisition rates from a pre-existent pool of HGT carriers are high. Plasmid transfer rates are likely increased at low AB doses^55^, but generally they are highly variable, and even though they are biased towards spread between clone-mates, there seems to be no obvious correlation between transfer rates and genetic distance of donors and recipients^56^. Hence, determining the relative importance of resistance evolution through HGT or chromosomal mutations is difficult, but for specific drugs like AMPs, for which spread of HGT resistance from the gut microbiota seems to be low^23^, the risk of treatment failure is mainly shaped by the beneficial mutations available to the population.

Most of our results assume that the pathogen population is completely susceptible to the drug at the onset of treatment. However, due to the fast growth and high mutation rates of bacteria, this is often not the case in real treatment scenarios and sometimes aiming for ‘mitigation’ (adaptive treatment) of the infection is more effective in reducing resistance evolution than trying to completely ‘eradicate’ the pathogen population (aggressive treatment). If multiple steps are necessary to obtain full resistance to the highest possible drug dose, we find that the threshold for choosing adaptive over aggressive treatment can be much lower than if only a single mutation were necessary (Fig. 4). In drug-free environments it is likely that for low benefits the majority of the population is more than one mutation away from full resistance, thereby providing a high competitive benefit paired with a low risk for resistance evolution. Even though determinations of resistant subpopulations are difficult in practice, this suggests that adaptive treatment is likely to be superior for many drugs, for which multi-step patterns are the most common resistance mechanism. Further, the assumptions in our model are not specific to bacterial populations or antimicrobials, which makes them more broadly applicable to other drug treatments, like cancer therapy^10^.

## Methods

### Literature review of costs and benefits of antimicrobial drug resistance mutations

We compounded a comprehensive set of experimental evolution studies (or reviews thereof) that measured both, fitness costs (usually growth rate reductions in the absence of drugs) and benefits (usually increases in MIC) of antibiotic (AB) or antimicrobial peptide (AMP) resistance mutations with the same set of experiments. We calculated costs as the arithmetic mean of 1-(relative fitness to wildtype) and the benefit as the geometric mean of MIC or IC_50_ increase relative to the wildtype. (Note: Chevereau et al. (2015)^25^ used IC_50_ instead of MIC but our calculation of IC_50_ and IC_90_ – which is likely very close to MIC - in their data gave a good correlation (R^2^=0.45, P<.001), which indicates that the benefits obtained from IC_50_ measurements are comparable to ones obtained from MIC measurements). As fitness measure we considered only the measurements done in the media that was also used for experimental evolution, even if growth was also measured in different environments. We obtained the type and average number of mutational events observed from supplemental data in most studies but there was generally no possibility to link any individual resistance mutation with a specific cost and benefit. Hence, we divided the overall costs and benefits by the average number of observed (adaptive) mutations (i.e. mutations that were not observed in control lines), assuming that each mutation provides a similar share to the overall magnitude. As most studies have a very low number of mutational events linked to resistance, this assumption is not expected to lead to strong biases. Overall, the results from all of the studies gave only a very weak positive linear correlation between the log(benefit) and cost of a mutational event (Fig. 1B). Mutations seem to be more likely to incur costs than benefits. This result is largely determined by the large data set from Spohn et al. (2019)^24^ which does not give a significant correlation between cost and benefit (Fig. S1), and the data points from Melnyk et al. (2015)^26^ which give a weak correlation. The dataset from Spohn et al. (2019)^24^ is the only one that fulfilled our criteria and directly compared AB and AMP mutational effects, which we summarize in Fig. S1.

### Pharmacodynamic model

We predicted the emergence of mutations in the population by using a pharmacodynamic model, which connects bacterial growth (or reductions thereof) to antimicrobial drug concentration via a Hill function (Fig. 1A)^1,5–7,11,57^. Benefits and costs were taken from the positive correlation that was observed with literature values (slope = 0.0087) - except for simulations testing the dependence of our results on this relationship, where we took a steeper correlation (slope = 0.0467) (Fig. S3). Model parameters other than benefit, cost and drug dose are taken from Yu *et al*. (2018)^11^ (Table S1). Bacteria can grow up to a certain carrying capacity and accumulate mutations a certain rate (we don’t allow for loss of mutations). Cells die at a low intrinsic rate, whereas death due to antimicrobials is given through the pharmacodynamic function and depends on the antimicrobial concentration as well as the MIC (see below). The change in each bacterial strain is then described as

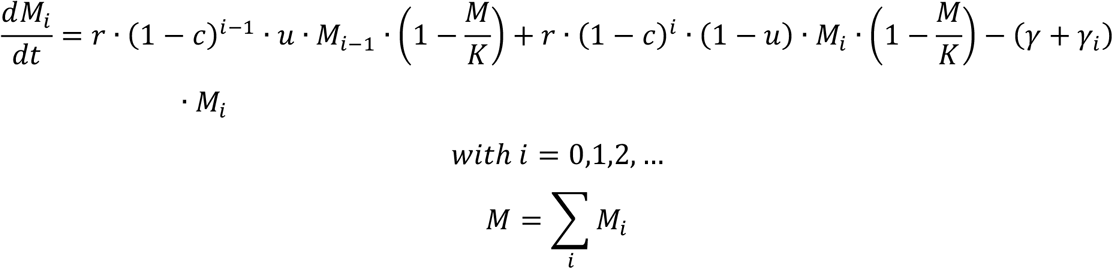

Here, M_i_ is the bacterial subpopulation carrying i mutations, r the growth rate (set to 1 in our simulations), c the cost of each mutation, u the mutation rate, K the carrying capacity of the system, *γ* the natural death rate and *γ*_*i*_ the death rate caused by drugs. *γ*_*i*_ is dependent on the properties of the antimicrobial applied and the benefit / cost conferred by each mutation. Hence, it is calculated from the maximal and minimal growth rates *ψ*_*max*_ and *ψ*_*min*_ (notice that *ψ*_*min*_ can be negative in the presence of drugs) the (time-dependent) concentration of the drug a, the MIC of the population (set to 1 in our simulations), the benefit b_i_ conferred by each mutation, and the sensitivity of the dose – growth relationship κ (the Hill coefficient or steepness of the curve):

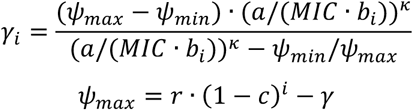

Accordingly, every sequential mutation decreases the maximum growth rate and decreases death due to the drug. Mutation rates were generally kept constant but considering higher mutation rates for mutations with lower benefits and costs did not change our results noticeably (Fig. S4). As benefits and costs are likely to vary, we also confirmed that our results are robust with regard to drawing benefits and costs of each mutation from a normal distribution. Similarly, we ran simulations with ‘peak PK’, where only the first mutational benefit / cost was fixed (i.e. deciding if a single- or multi-step pattern was necessary) and the other mutations were sampled from the whole range of benefits and costs obtained from the literature, independently of each other (Fig. S13). We also tested the impact of the mutation rate on our results and increased the mutation rate for small benefit / cost mutations proportional to the number of mutational steps that were needed to gain resistance. Based on previous experimental and theoretical work^11,53,58^, we defined two different antimicrobial drug classes by using two parameter sets: for the AB class the mutation rate was 3*10^−6^, κ was 1.5 and *ψ*_*min*_ was -5; whereas for the AMP class the mutation rate was 10^− 6^, κ was 5 and *ψ*_*min*_ was -50 ^11^.

### Pharmacokinetic functions

In our simulations we used three different PK functions to evaluate resistance evolution dynamics. ‘Peak PK’ describes the intake of a drug with a certain frequency τ, which is absorbed instantaneously and then decays exponentially at rate k^11^:

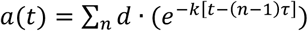

with n = 1,2,… the number of times the treatment dose d is applied. For ‘constant PK’ the drug concentration is independent of time and simplifies to a=d, whereas for ‘ramp PK’, the drug concentration increases linearly over a time k2cmax (hence the rate of drug concentration increase is given as: d/k2cmax) and then stays constant for the rest of the treatment period. The value for k2cmax used for most simulations (48h) was taken from literature and describes an example of AMP production timing during a natural immune response ^39^.

### Implementation and simulation

The model was implemented in R, using the package *deSolve*^59^ for deterministic analysis and the package *adaptivetau*^60^ for stochastic implementations via the Gillespie algorithm. We calculated treatment failure probability as the frequency of runs with surviving bacteria at the end of the treatment period (200h).

Mutational diversity was calculated using the Shannon index, which takes into account the richness and evenness of the distribution of mutant subpopulations, and was averaged over the treatment period. The values for treatment failure and diversity were then averaged over the whole multi- or single-step area (triangles shown in Fig. 2) in order to compare different treatment strategies.

The number of mutations that escaped drift (mutational radius) in a single run was taken as the subpopulation with the highest number of mutations that reached at least 10% of the whole population (with the whole population being at least as big as the starting population, i.e. 10^6, to avoid counting stochastic survival at very low population densities) and then averaged over all runs.

The difference between different antimicrobial classes was obtained by subtracting the corresponding values after every simulation of an AMP treatment from the one obtained in a simulation for an AB treatment and plotting the individual resulting differences (for 500 simulations) as well as the density via violin plots.

The described code will be made available as an R package.

### Selection coefficient analysis

Selection coefficients were calculated under the assumption that the sensitive population is very small compared to the carrying capacity, which means that we can neglect the logistic growth term in our calculations. As the results were very similar to assuming a population at the carrying capacity, we will focus on the selection coefficients with a small starting population. Selection coefficients were determined through eigenvalues obtained from the Jacobi matrix given by 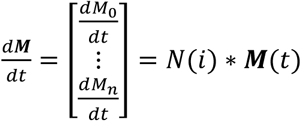:

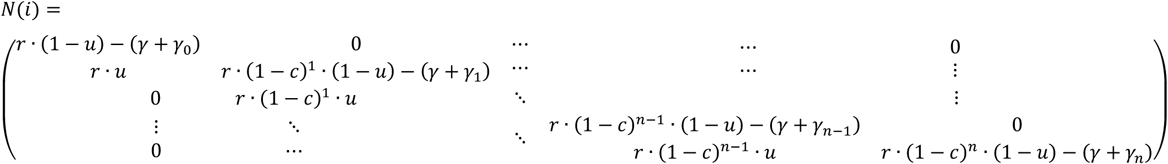

The Eigenvalues of M(A) are its diagonal entries, which correspond to the net growth of each population. We calculated selection coefficients as the difference in growth rates between bacteria with i mutations and bacteria with i-1 mutations (i.e. the difference between their Eigenvalues):

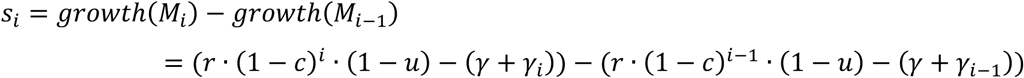

The difference between the parameter sets for the two antimicrobial classes used here lies in the higher mutation rate, lower κ and higher *ψ*_*min*_ for AB treatments ^11^. Hence, we investigated the importance of the two PD parameters κ and *ψ*_*min*_ by calculating the selection coefficients using the AMP parameter set and swapping either κ or *ψ*_*min*_ with that of the AB parameter set.

More generally, we can consider the Jacobi matrix for the resistant populations invading at the mutant-free equilibrium:

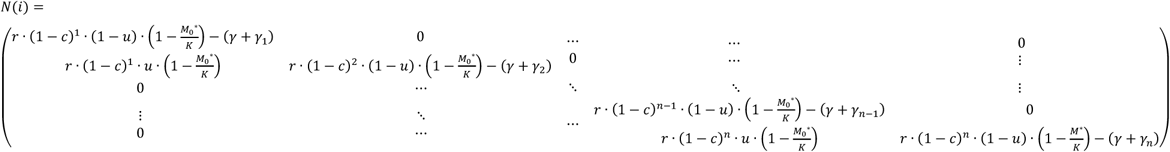

The criteria for invasion of a mutant into the susceptible population is then that the eigenvalue of the mutant has to be bigger than zero, i.e.

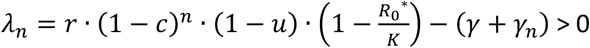

Inserting the mutant-free equilibrium 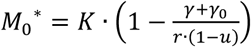 yields:

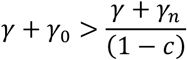

Which means that bacterial cells with n mutations can invade if the death rate of the sensitive strain is higher than the death rate of the mutant normalized by the cost of the mutation(s).

### Horizontal gene transfer (HGT)

We added HGT to the model by allowing for an additional resistance gene (with benefit b_p_ and cost c_p_) to be acquired, which gives resistance in a single step. Hence, the benefit b_p_ and the corresponding cost c_p_ were adjusted according to the (maximum) drug dose applied. This gene can be acquired at a low rate *α* from the environment, or at a density dependent rate *β*, which we assumed to be on the same order of magnitude as the mutation rate ^61^. The transferred gene can be acquired by bacterial populations with or without mutations and cells containing the transferred resistance can still acquire further mutations (but not further HGT resistance). The equations were modified as follows:

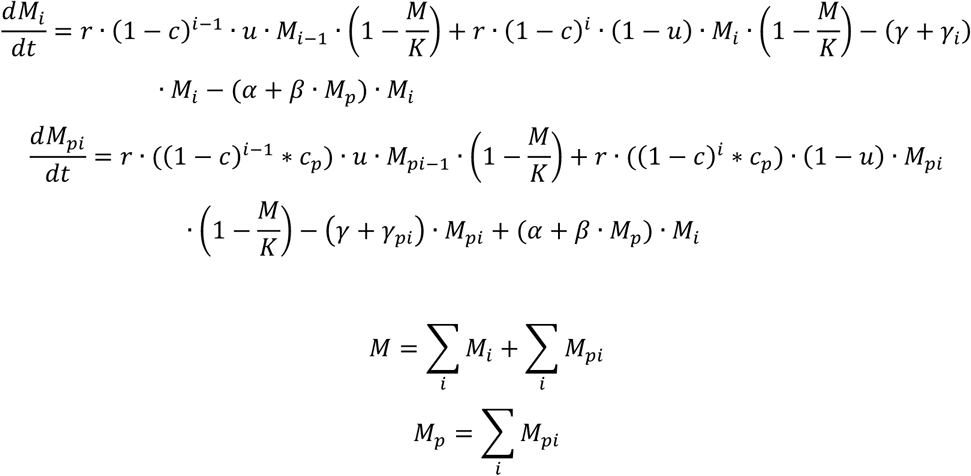

Here, M_pi_ is the bacterial subpopulation carrying the HGT gene and i mutations, M_p_ the total number of HGT subpopulations and M the total number of all bacterial populations. Relative population frequencies were calculated at the end of the treatment period by dividing the cell number of each subpopulation through the whole population size.

### Adaptive treatment

In adaptive treatment the goal is not to eradicate the bacterial population entirely but to adjust the treatment dose continuously in order to keep the pathogen level below a certain upper limit. Hansen, Woods and Read (2017)^18^ calculated the threshold of resistant cells that are necessary at the beginning of the treatment for adaptive treatment to outperform aggressive treatment (which constitutes giving the full dose right away), which is based on the idea that sensitive cells provide a risk for becoming resistant through mutation and a benefit through growth competition with the resistant cells at the same time. Hansen, Woods and Read (2017)^18^ only considered one mutation to resistance, which means that their risk subpopulation and competitior subpopulation was the same. If we consider however sequential mutational steps, then the risk population only consists of the subpopulation one mutation away from full resistance (which will be the m-th mutation), whereas the competitor population for the fully resistant strain contains all (partially) sensitive bacteria (i.e. including mutant strains, which are not fully resistant to the highest possible treatment dose). Therefore, the threshold of resistant bacteria is given by (compare to (4) in Hansen, Woods and Read, 2017):

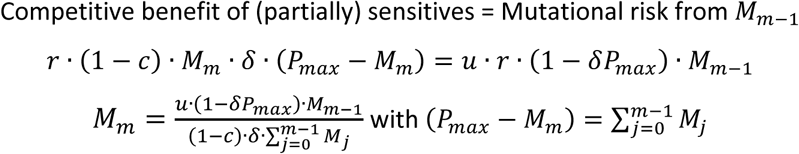

Here, *δ* describes the strength of competition and *P*_*max*_ the upper limit of acceptable pathogen burden. This leads to a quadratic equation for the subpopulation with m mutations, *M*_*m*_, which we used to calculate how the resistant population threshold for adaptive treatment (i.e. the initial density *M*_*m*0_ above which adaptive treatment is more favorable) differs between multi- and single-step resistance patterns (Fig. 4).

We implemented adaptive treatment in our model by setting an upper bound of acceptable pathogen cells, and adjusting the treatment dose in order to keep the pathogen load at or below this threshold but the subpopulations of at least partially sensitive cells as big as possible (Fig. S9). These (partially) sensitive cells serve as competitors for the resistant strain which carries a mutational growth cost and can be outcompeted at low drug doses^18^. At the same time subpopulations that are one step away from the resistant population provide a risk population as they are likely to gain resistance.

For simulations of adaptive and aggressive treatment, we started from a population with neutral heterogeneity, meaning that we calculated the staedy state number of cells with a specific number of mutations given a certain cost (and benefit) in the absence of drug selection. As we want to compare the time difference to treatment failure between adaptive and aggressive treatment for single- and multi-step patterns, we initially add to this ‘neutral population’, the number of resistant cells necessary to make adaptive treatment optimal. The drug dose in adaptive treatments was then adjusted to keep the number of pathogens below the acceptable burden and by increasing the dose according to the increase in partially resistant subpopulations. The time of treatment failure was determined as the time where the total pathogen population crossed 10^8^ CFUs. We compared adaptive and aggressive treatment by dividing the time to treatment failure obtained from the adaptive strategy by the one obtained with the aggressive strategy, yielding the fold difference in treatment success duration.

## Acknowledgements

We thank D. Baeder, S. Bonhoeffer, S. Lehtinen and H. Alexander for useful discussions and comments on the manuscript. This work was supported by a grant from the Volkswagen Foundation (grant nr. 96517).

## Supplementary Figures and Tables

**Table S1.** Parameters used in the PD model.

| Parameters | Value | Unit | Description |
| --- | --- | --- | --- |
| $r$ | 1 | $h^{-1}$ | Replication rate |
| $b$ | 2-100 | xMIC | Benefit per mutation |
| $c$ | 0.008-0.2 | - | Cost per mutation |
| $u$ | $3 \cdot 10^{-6}$,<br>$1 \cdot 10^{-6}$ | - | Mutation rate (AB, AMP) |
| $K$ | $10^9$ | CFU | Carrying capacity |
| $\gamma$ | 0.01 | $h^{-1}$ | Death rate |
| $\psi_{min}$ | -5, -50 | $h^{-1}$ | Min. growth rate (AB, AMP) |
| $a$ | 0.1-100 | xMIC | Drug concentration |
| $k_a$ | 0.5 | $h^{-1}$ | Drug absorption rate |
| $k$ | 0.1 | $h^{-1}$ | Drug decay rate |
| $\kappa$ | 1.5, 5 | - | Steepness of the PD curve (AB, AMP) |
| $\tau$ | 1/24 | $h^{-1}$ | Dose frequency |
| $K2cmax$ | 12-96 | h | Time till max. dose |
| $M_i$ | $0-10^9$ | CFU | Population with $i$ mutations |
| $M_{pi}$ | $0-10^9$ | CFU | Population with a plasmid and $i$ mutations |
| $b_p$ | 0.6-24 | xMIC | Benefit of the plasmid |
| $c_p$ | 0.03-0.1 | - | Cost of the plasmid |
| $\alpha$ | $3 \cdot 10^{-10}$ | $h^{-1}$ | Plasmid acquisition rate from the environment |
| $\beta$ | $3 \cdot 10^{-8}$ | $h^{-1}$ | Plasmid acquisition rate from $M_{pi}$ carriers |
| $P_{max}$ | $10^5$ | CFU | Max. acceptable pathogen burden |

**Figure S1.**
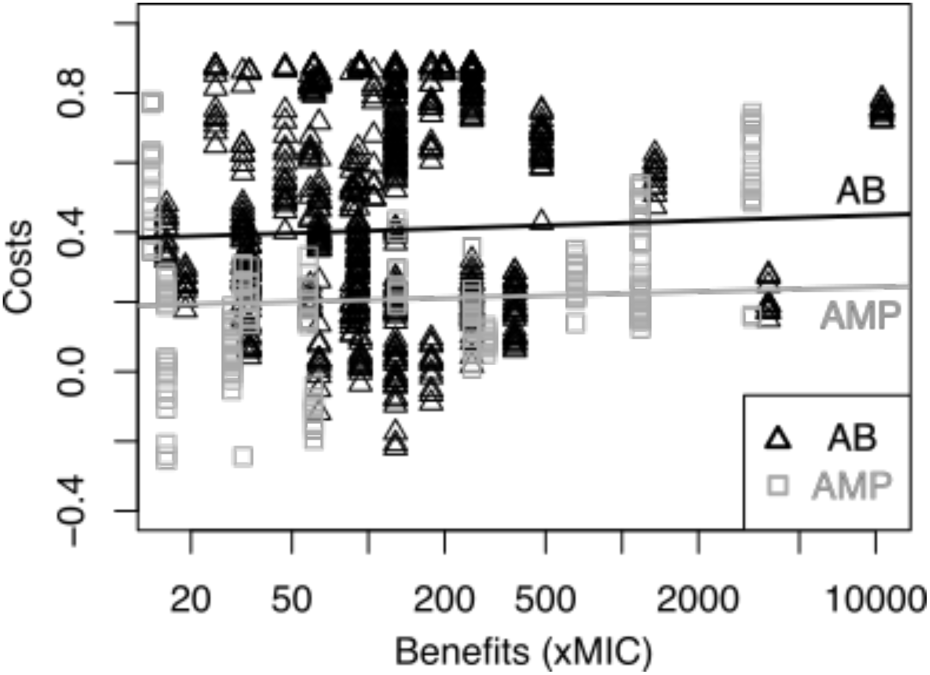
Direct comparison of mutational benefits and costs between AMPs and ABs from Spohn *et al*., 2019. Cost (relative growth reduction) and benefit (MIC increase) of mutations at the end of the evolution experiments with AMPs (grey rectangles) or ABs (black triangles) from Spohn *et al*., 2019 are shown. We find only a weak negative correlation (obtained through linear regression) between the costs and (log) benefit. This could be due to the fact that in this study five mutations were present in one genome on average and compensatory mutations could confound the actual costs of resistance mutations.

**Figure S2.**
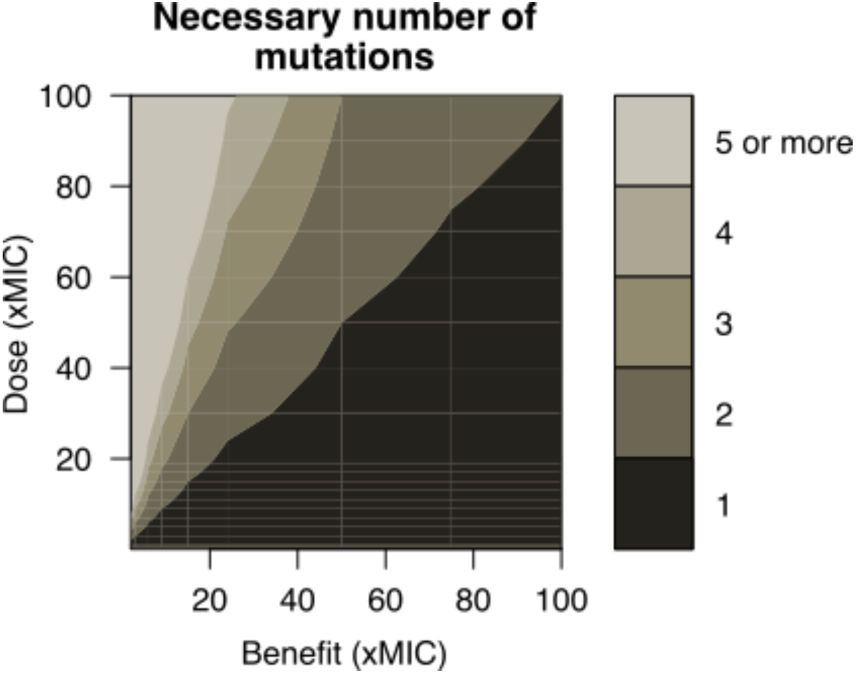
Number of mutations necessary for resistance. The combination of a given drug dose (xMIC) and the benefit per mutation (xMIC) determines the number of mutations necessary for bacterial growth in our PD model: single-step evolution occurs when only 1 mutation is needed (dark grey area), whereas multi-step resistance requires 2 or more mutations (lighter grey areas).

**Figure S3.**
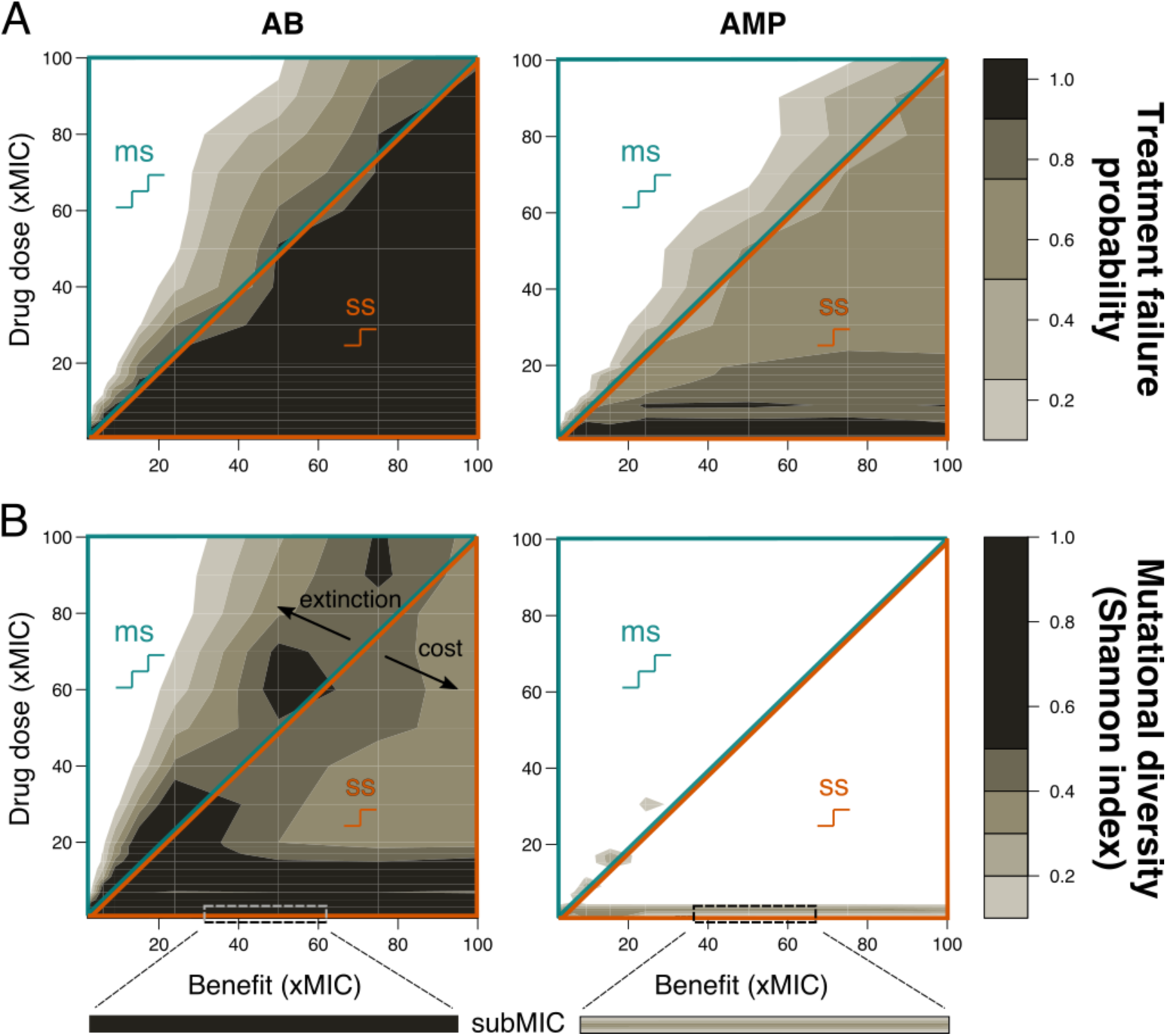
Resistance evolution with single- and multi-step patterns for peak PK with steeper correlation between cost and log(benefit). A) Treatment failure probability and B) mutational diversity are shown for two different antimicrobial classes (ABs – left, AMPs – right) for different combinations of mutational benefits (xMIC) and drug doses (xMIC). The diagonal line separates single-step (ss, lower orange triangle) from multi-step (ms, upper blue triangle) resistance. A representative example of subMIC mutational diversity is shown magnified below the plots in B).

**Figure S4.**
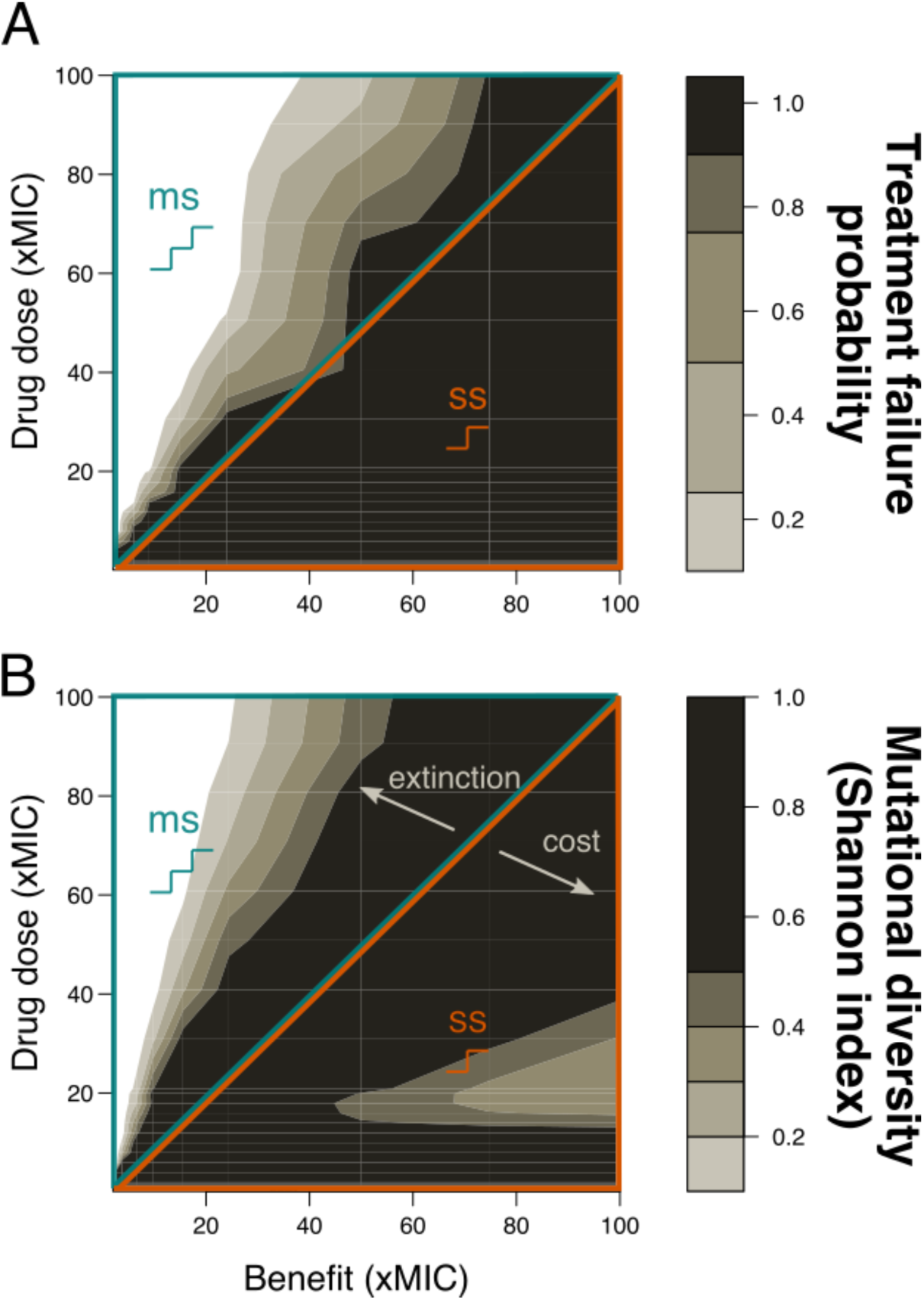
Resistance evolution with increased mutation rates (proportional to the number of mutations required for resistance). A) Treatment failure probability and B) mutational diversity are shown for AB treatments with peak PK for different combinations of mutational benefits (xMIC) and drug doses (xMIC). The diagonal line separates single-step (ss, lower orange triangle) from multi-step (ms, upper blue triangle) resistance Mutation rates were increased for lower-benefit mutations, which resulted in increased mutational diversity, but not higher treatment failure.

**Figure S5.**
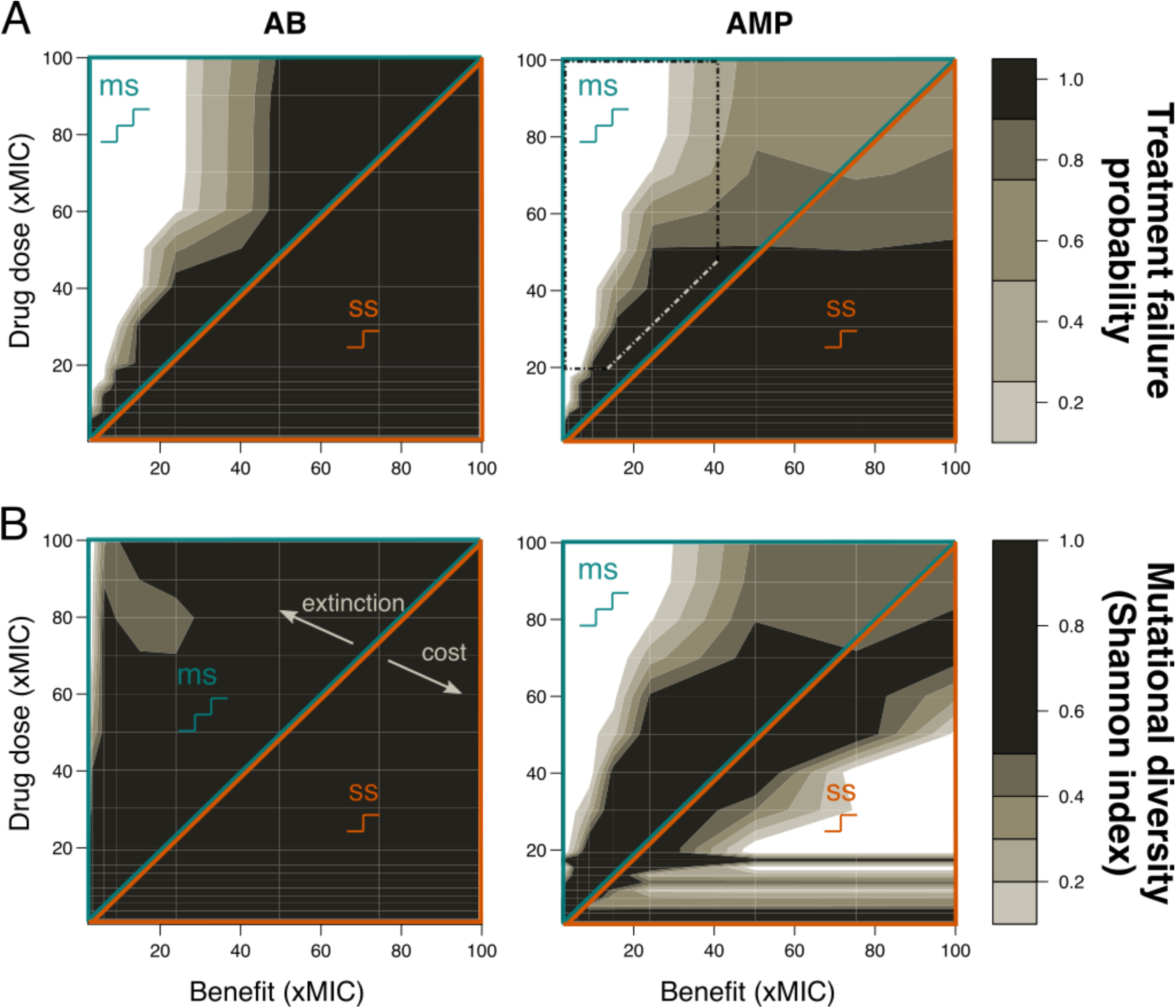
Resistance evolution with single- and multi-step patterns for ramp PK. A) Treatment failure probability and B) mutational diversity are shown for two different antimicrobial classes (ABs – left, AMPs – right) for different combinations of mutational benefits (xMIC) and drug doses (xMIC). The diagonal line separates single-step (ss, lower orange triangle) from multi-step (ms, upper blue triangle). The dotted rectangle shows the area in which AMPs lead to higher treatment failure probabilities than ABs.

**Figure S6.**
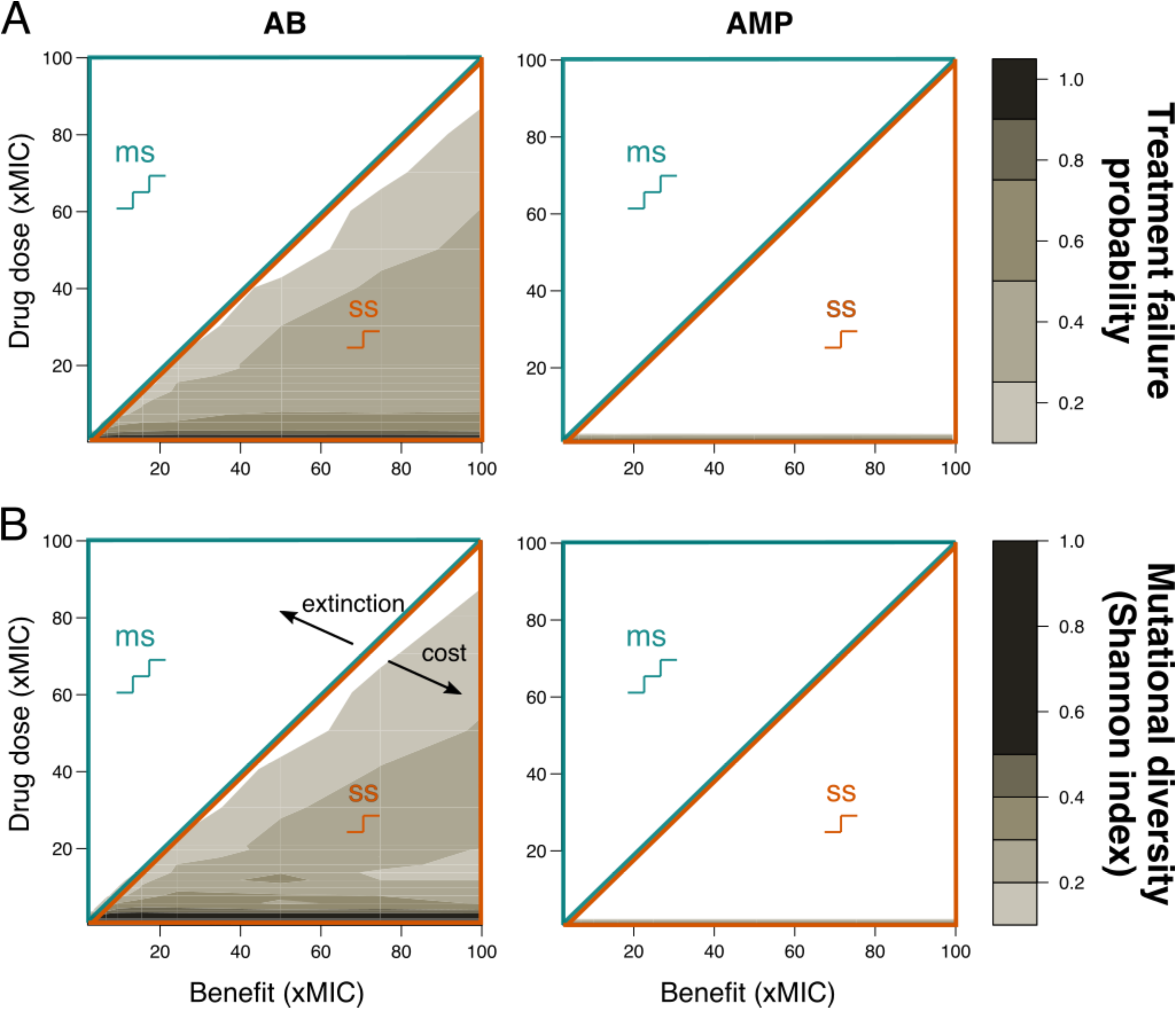
Resistance evolution with single- and multi-step patterns for constant PK. A) Treatment failure probability and B) mutational diversity are shown for two different antimicrobial classes (ABs – left, AMPs – right) for different combinations of mutational benefits (xMIC) and drug doses (xMIC). The diagonal line separates single-step (ss, lower orange triangle) from multi-step (ms, upper blue triangle) resistance

**Figure S7.**
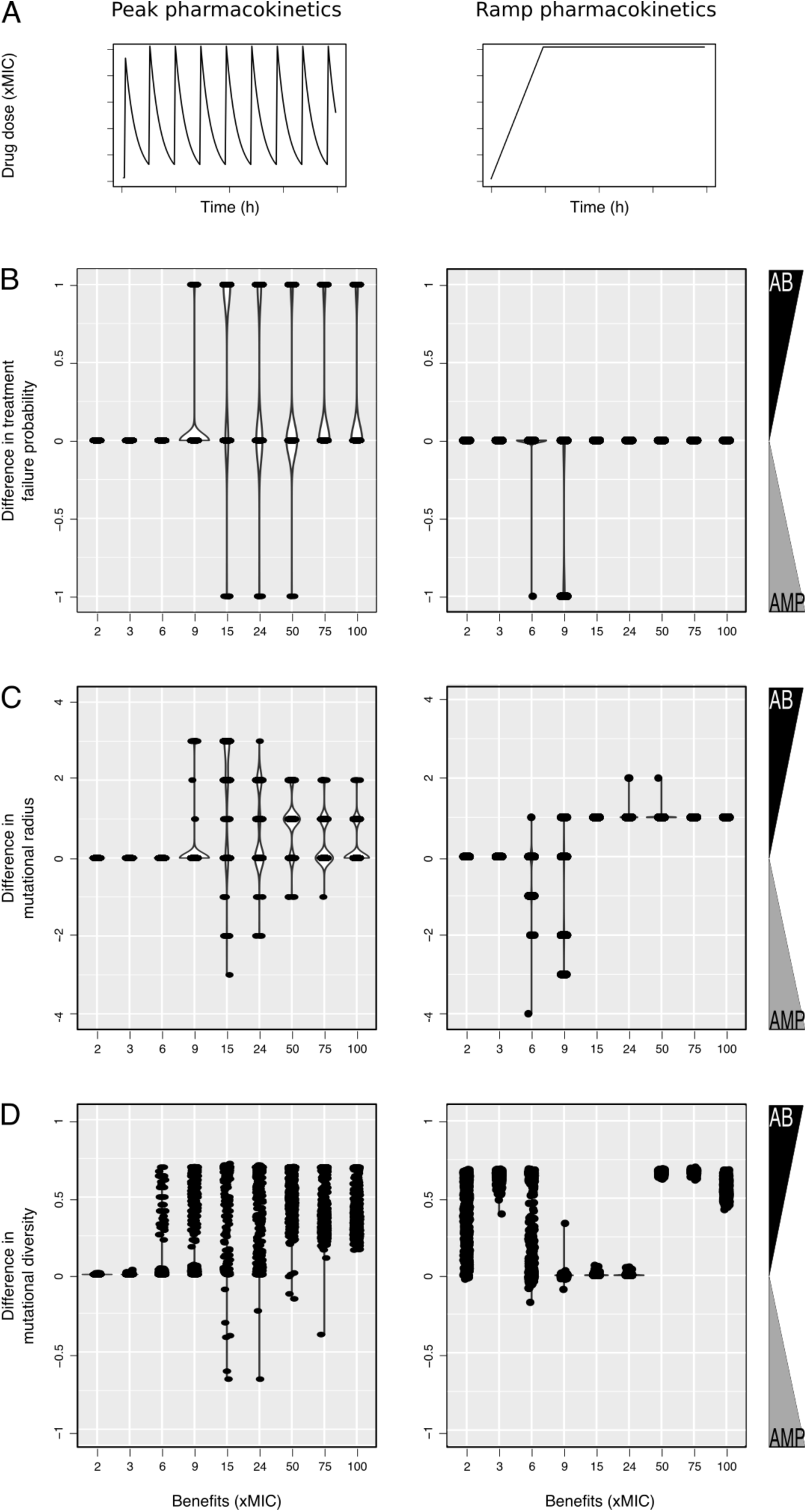
Comparison of resistance evolution with AB and AMP treatments using peak or ramp PK (A). Violin plots show the densities, as well as the individual simulation results, obtained from calculating the difference between AB and AMP treatments regarding B) treatment failure, C) number of mutations surviving genetic drift, and D) mutational diversity after every simulation with positive values showing higher incidence in AB treatments and negative values, higher incidence in AMP treatments. Simulations were run with a ramp PK increase of 48h at 20xMIC drug dose for various benefits as shown.

**Figure S8.**
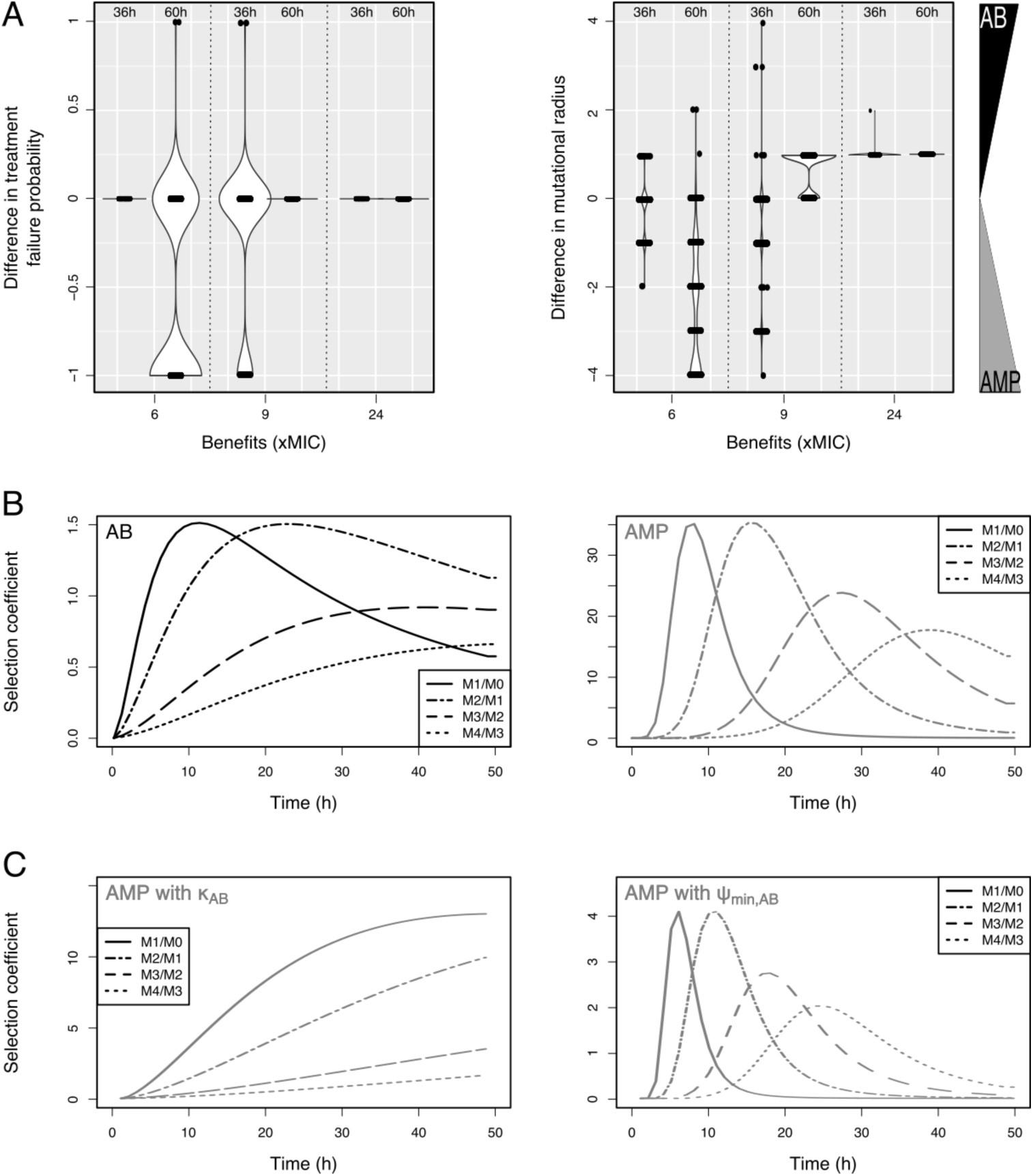
Selection coefficient analysis. A) Violin plots give the density and individual points for the difference between AB and AMP treatment failure probability calculated after every simulation run with a ramp PK increase of 36h or 60h at 20xMIC drug dose for various mutational benefits as shown. Positive values showing higher incidence in AB treatments and negative values, higher incidence in AMP treatments. B) Selection coefficients (see Methods for calculation) for a (final) drug dose of 20xMIC, a ramp time of 48h and a benefit per mutation of 2xMIC are shown for AB (left) or AMP (right) treatments. Solid lines give the selection of the first mutant over the wildtype, dash-dotted lines selection of the second over the first mutant, dashed lines selection of the third over the second mutant and dotted lines selection of the fourth over the third mutant. C) Selection coefficients for the same conditions as in B) with AMP treatments but the PD parameter κ (left) or *ψ*_*min*_(right) are swapped with the ones for AB treatments. Whereas κ changes the selection coefficients in a qualitative manner, *ψ*_*min*_ does so in a quantitative manner.

**Figure S9.**
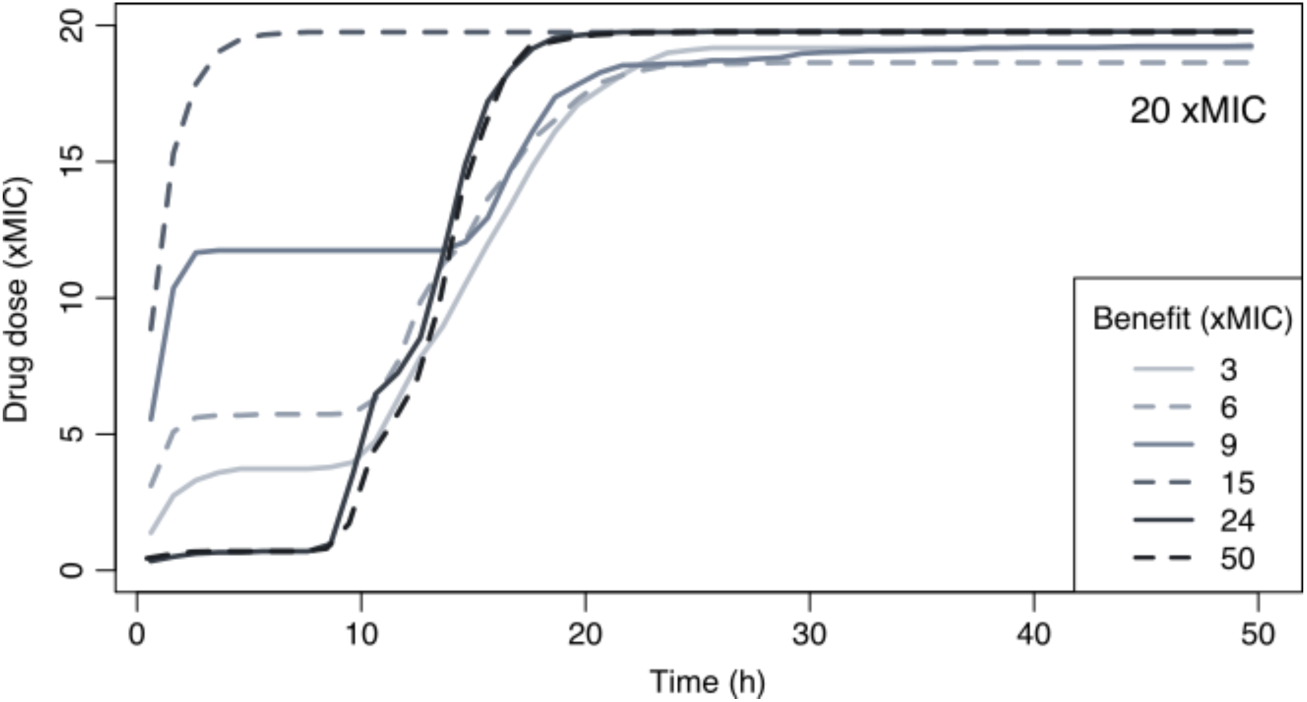
Optimal PK kinetics for adaptive treatment. Shown are the optimal PKs (xMIC) over time(h) if drug doses are adapted to keep the bacterial population below a certain maximum but the number of competitors as high as possible. The maximum drug dose was set as 20xMIC and the mutational benefit was varied between 3 and 50 (xMIC).

**Figure S10.**
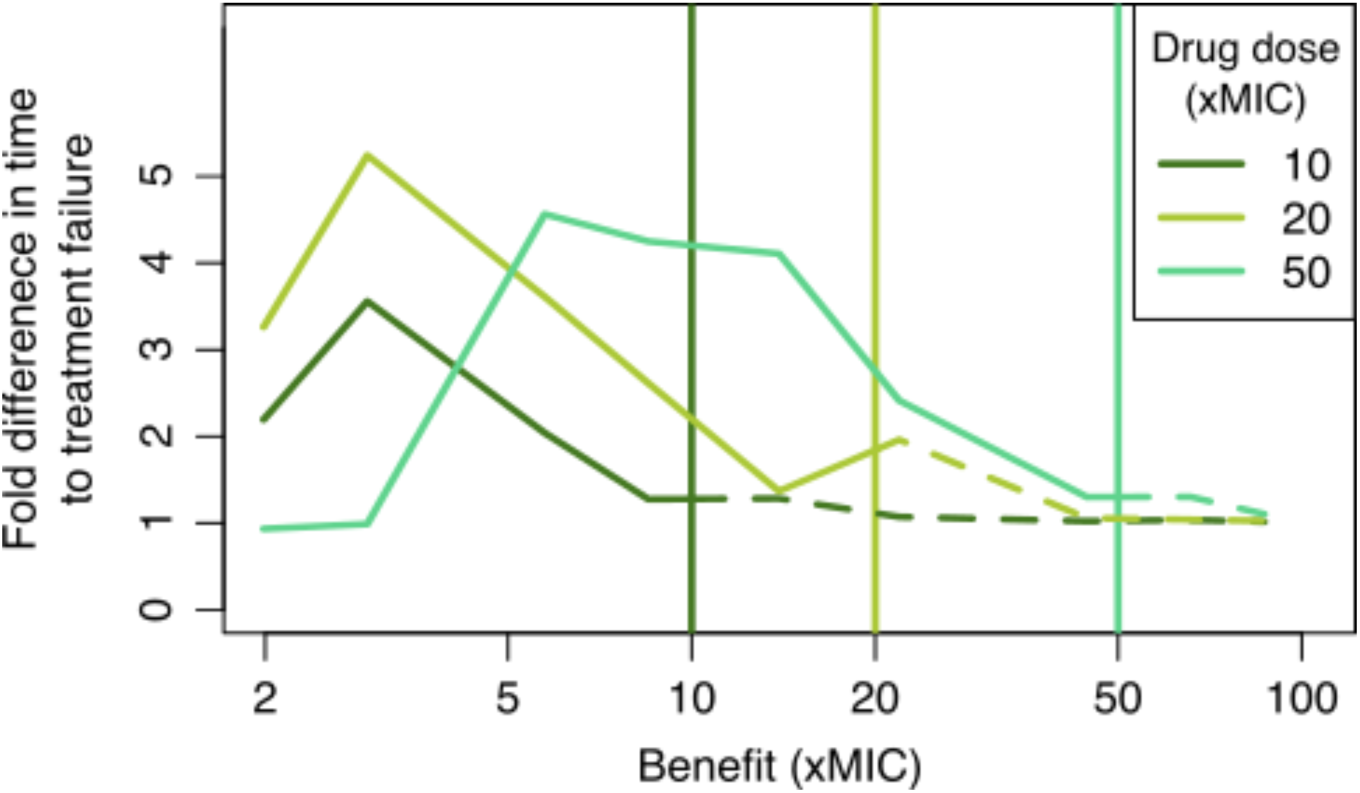
Differences in time to treatment failure are more pronounced with multi-step patterns. Shown is the fold difference between aggressive (peak PK) and adaptive treatment in the time needed for the pathogens to escape treatment at a specific maximum drug dose (xMIC) for a range of mutational benefits (xMIC), starting with a population containing enough mutants to favor adaptive treatment. Vertical lines show the respective thresholds dividing multi-step from single-step resistance patterns (single-step patterns are shown in dashed lines). Particularly for multi-step resistance evolution, adaptive treatment can lead to much longer time periods before treatment failure, but we also find a trade-off: if the pathway to resistance involves too many steps than keeping the competitive population alive requires very low doses, whereas the few resistant bacteria do not pay enough growth cost to delay treatment failure longer than with aggressive treatment.

**Figure S11.**
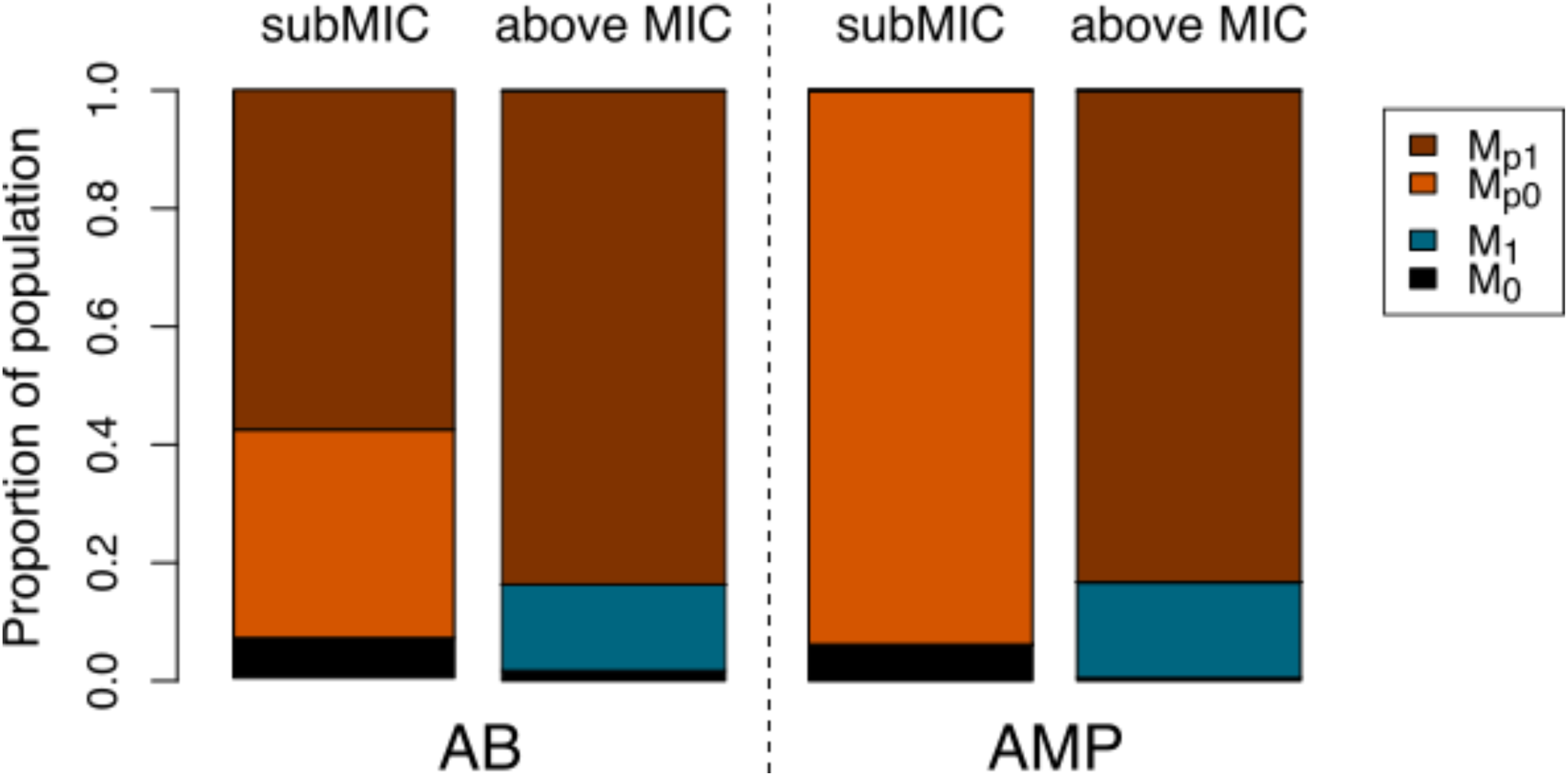
Relative population frequencies with horizontal gene transfer (HGT). Shown are representative examples of population frequencies at the end of drug treatments (AB or AMP) at subMIC or above MIC levels. Black indicates the wildtype (completely sensitive), blue the population fraction with 1 mutation and orange (brown) the population fraction with a plasmid and 0 (1) mutation.

**Figure S12.**
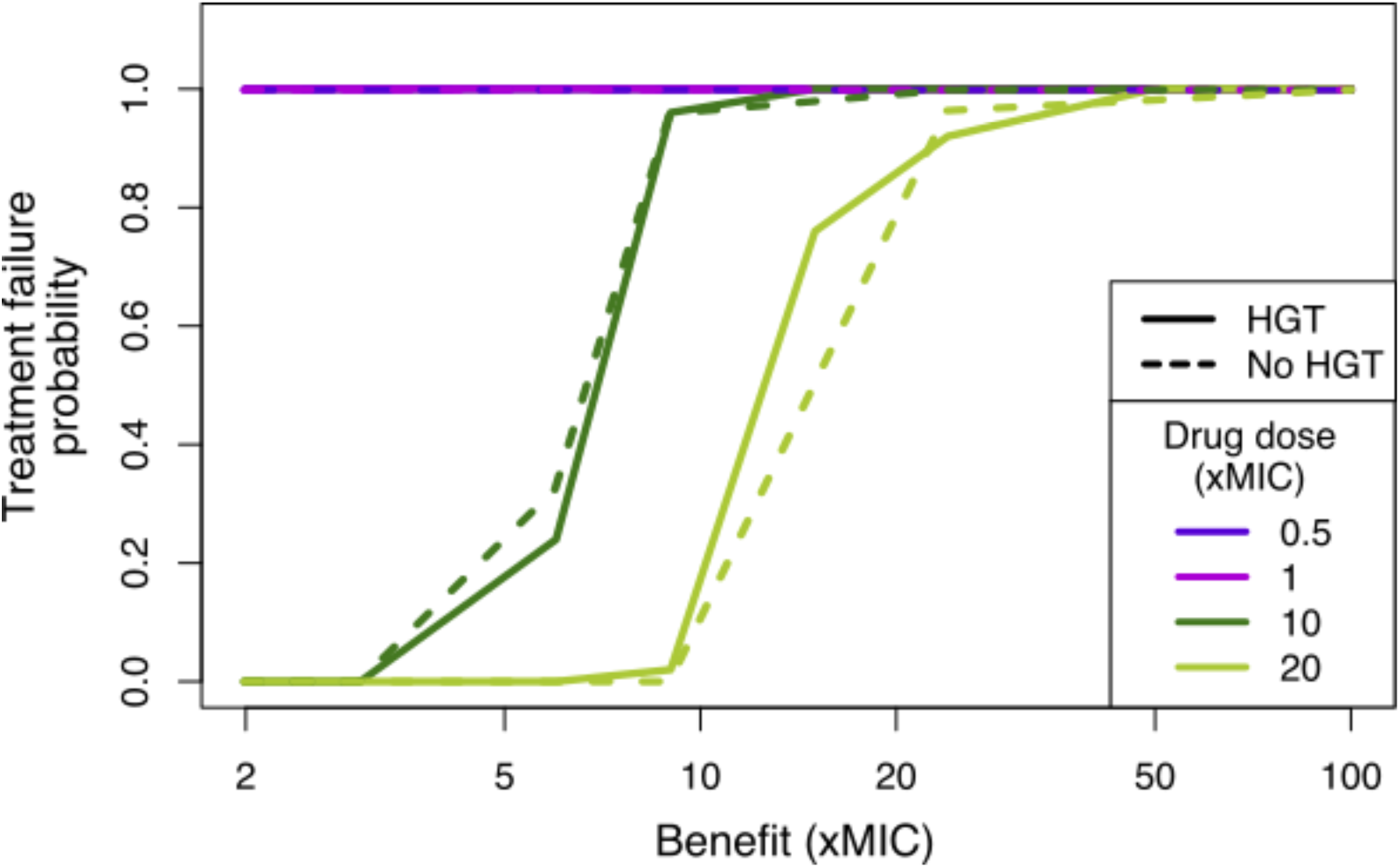
Treatment failure is similar with and without HGT. The probability of treatment failure with PK kinetics are shown for four AB doses (xMIC) in simulations with (solid lines) or without (dashed lines) HGT over a range of mutational benefits (xMIC).

**Figure S13.**
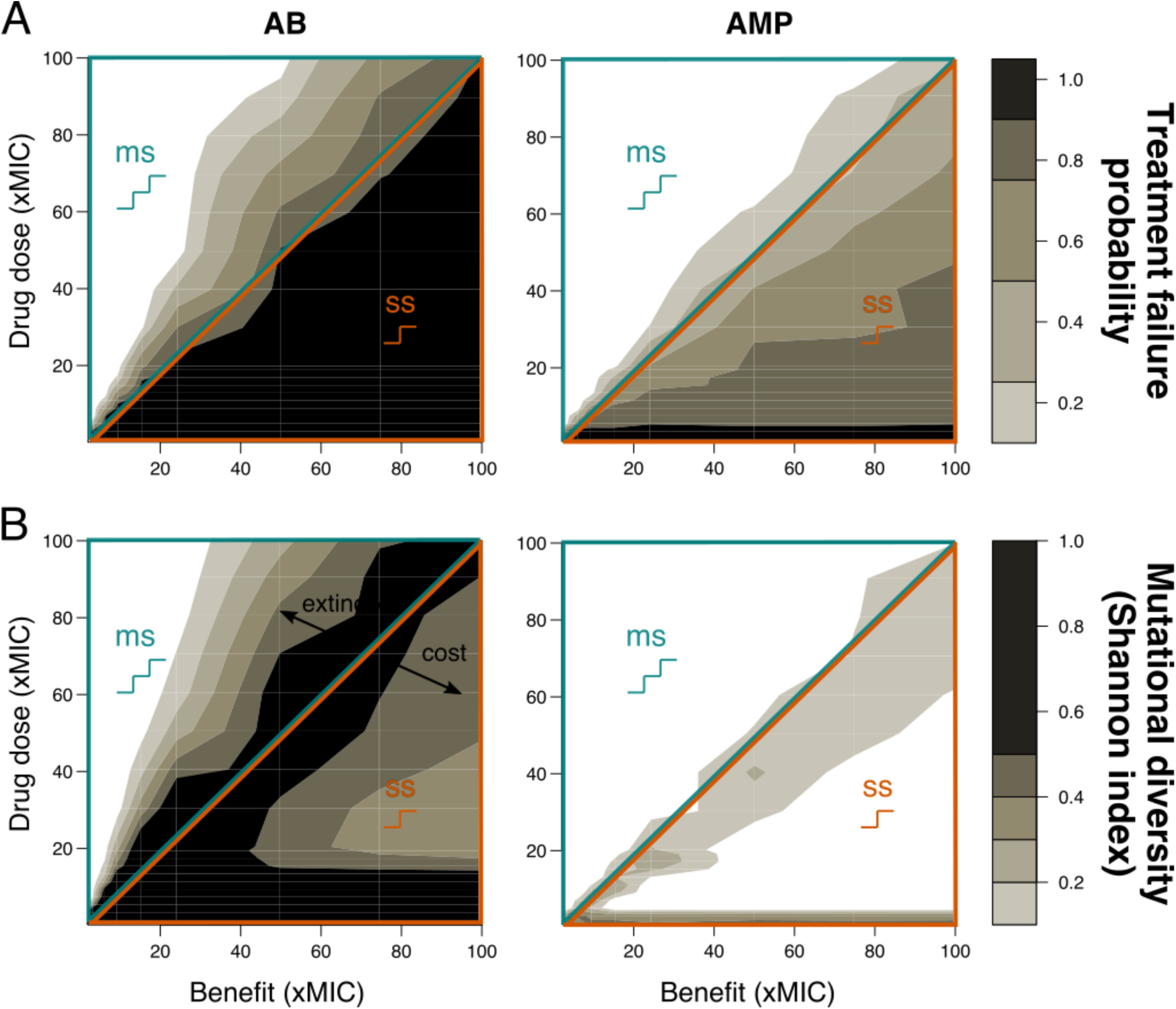
Resistance evolution with random mutational benefit and cost. Benefit and cost of the first mutation were pre-determined (x-axis), determining single-step (ss) or multi-step (ms) resistance patterns, but further mutational benefits and costs were drawn randomly and independently. A) Treatment failure probability and B) mutational diversity are shown for two different antimicrobial classes (ABs – left, AMPs – right) for different combinations of mutational benefits (xMIC) and drug doses (xMIC). The diagonal line separates single-step (ss, lower orange triangle) from multi-step (ms, upper blue triangle) resistance

